# Blebs Promote Cell Survival by Assembling Oncogenic Signaling Hubs

**DOI:** 10.1101/2021.04.23.441200

**Authors:** Andrew D. Weems, Erik S. Welf, Meghan K. Driscoll, Felix Zhou, Hanieh Mazloom-Farsibaf, Bo-Jui Chang, Vasanth S. Murali, Gabriel M. Gihana, Byron G. Weiss, Joseph Chi, Divya Rajendran, Kevin M. Dean, Reto Fiolka, Gaudenz Danuser

## Abstract

Most human cells require anchorage for survival. Cell-substrate adhesion activates diverse signaling pathways, without which cells undergo anoikis – a form of programmed cell death^1^. Acquisition of anoikis resistance is a pivotal step in cancer disease progression, as metastasizing cancer cells often lose firm attachment to surrounding tissue^2–5^. In these poorly attached states, cells adopt rounded morphologies and form small hemispherical plasma membrane protrusions called blebs^6–13^. Bleb function has been thoroughly investigated in the context of amoeboid migration but is far less examined in other scenarios^14–19^. Here we show by quantitative subcellular 3D imaging and manipulation of cell morphological states that blebbing triggers the formation of plasma membrane-proximal signaling hubs that confer anoikis resistance. Specifically, we discovered in melanoma cells that blebbing generates plasma membrane contours, which recruit curvature-sensing septin proteins as scaffolds for constitutively active mutant NRAS and effectors. These signaling hubs activate ERK and PI3K – canonical promoters of pro-survival pathways. Inhibition of blebs or septins has little effect on the survival of well-adhered cells, but in detached cells causes NRAS mislocalization, reduced MAPK and PI3K activity, and ultimately, death. This unveils an unanticipated morphological requirement for mutant NRAS to operate as an effective oncoprotein. Moreover, we find that though some BRAF mutant melanoma do not rely on this survival pathway in a basal state, BRAF/MEK inhibition strongly sensitizes them to both bleb and septin inhibition. Importantly, we demonstrate that fibroblasts engineered to sustain blebbing acquire the same anoikis resistance as cancer cells even without harboring oncogenic mutations. These data define a role for blebs as potent signaling organelles capable of integrating myriad cellular information flows into concerted cellular responses, in this case granting robust anoikis resistance.

## Main

Attachment-dependent survival is a fundamental aspect of metazoan cell physiology. Near universally, points of attachment – either to the extracellular matrix or other cells – generate pro-survival signals, often through the assembly of ultrastructural cellular architecture such as focal adhesions^20,21^ and adherens junctions ^22–24^. These adhesive structures operate as signaling hubs by controlling local concentrations of signaling factors, protein scaffolds, and effector proteins^25^. If deprived of attachments, most cells will undergo a detachment-induced form of programmed cell death called anoikis^1^. Acquisition of molecular strategies that confer anoikis resistance is a critical step in oncogenesis^2–4^. Though many mechanisms of anoikis-resistance in cancer are known, understanding of this phenomenon is far from complete^5^.

Upon detachment, most metazoan cell types become rounded and display dynamic surface blebs^7–13^ (**Movie 1**). Blebs are hemispherical, pressure-driven cellular protrusions that occur when portions of the cell membrane decouple from the actomyosin cortex^6^. Though blebbing famously occurs during apoptosis^26–28^, a second class of “dynamic blebs” are found in healthy cells with intact cortices and associated with cellular detachment, mitosis, and amoeboid motility^6,28^. Detached nonmalignant cells maintain dynamic blebbing for only 1-2 hours, later undergoing anoikis if attachment is not re-established^11,12,29^. In contrast, cancer cells, such as those forming melanoma, are highly anoikis-resistant and able to indefinitely sustain rounded blebby morphologies in no- and low-attachment environments, often adopting this amoeboid-associated phenotype both *in vitro* and *in vivo* – notably at the invasive front of tumors^14–19,30^. This increased blebbiness has recently been associated with increased metastatic potential in melanoma and prostate cancer^14,31^, suggesting the morphotype might be an indicator of disease aggressiveness.

The correlation between blebby morphologies and metastasis has largely been attributed to the role of blebs in cell motility^14–19^, although recent work has begun to suggest that blebbiness might also convey a survival advantage^30^. This would agree with our previous finding that signaling factors associated with cell survival tend to be concentrated in the vicinity of melanoma blebs, including KRAS and active PI3K^14,32^. Similar bleb-localized enrichments of ERK1/2 have also been observed^33^. Moreover, inhibition of blebbing causes a sharp reduction in local PI3K activity, raising the possibility that blebbing might directly enable this signaling^14^. Taken together, this information motivated an investigation of blebbing as a potential contributor to anoikis resistance in melanoma.

### Bleb inhibition disrupts anoikis resistance in melanoma cells

We first inhibited bleb formation using wheat germ agglutinin (WGA), a lectin that reduces plasma membrane deformability by crosslinking components of the glycocalyx^6^. Using MV3 melanoma cells, we confirmed by morphological motif analysis^32^ WGA’s effects as a dosage dependent inhibitor of bleb density on single cell surfaces (**Fig 1A**). To investigate putative bleb function as a driver of anchorage-independent cell survival we designed assays that consistently minimized both cell-substrate and cell-cell attachment over a period of 24 hours. We accomplished this goal by seeding cells at very low density (approx. 100 cells/ml) in non-adherent culture dishes under constant gentle nutation, which we found necessary to prevent cell aggregation. For controls we grew cells at identical concentrations in adherent variants of the same culture dishes. Together, this generated pairs of ‘detached’ and ‘adhered’ cell populations in paired same-day experiments.

**Figure 1:**
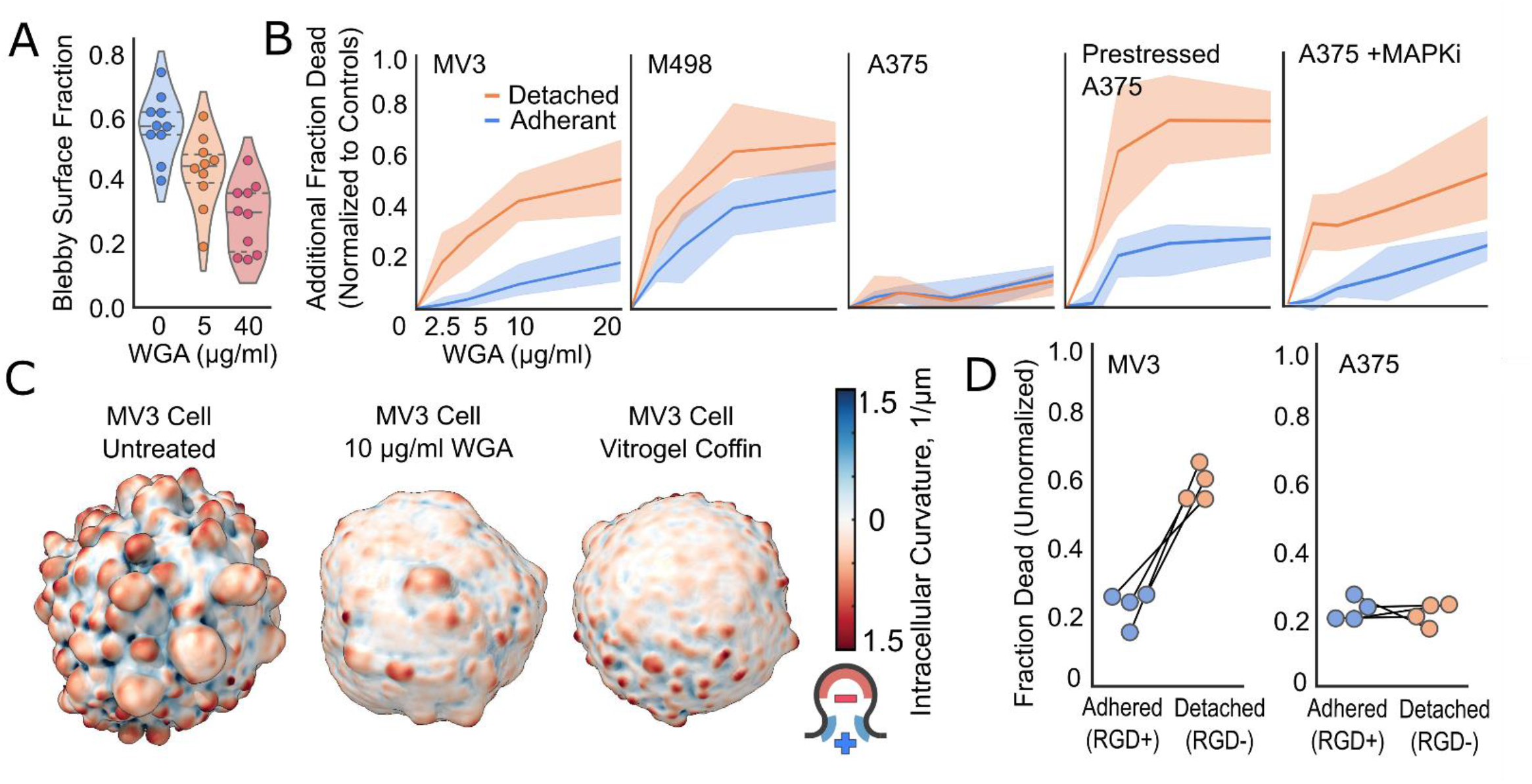
Bleb inhibition disrupts anoikis resistance in melanoma cells. **(A)** Fraction of cell surface comprised of blebs for MV3 cells treated with different concentrations of WGA. Dashed lines separate quartiles. Dots represent individual cells. **(B)** Cell death as a function of WGA treatment at different doses for adhered and detached melanoma cells. Cells grown for 24 hours and assayed for cell death using ethidium homodimer staining. All treatment groups grown and assayed in simultaneous paired experiments, which were performed three times for each cell line (See Fig. S1 for individual experiments). Replicate data were normalized by subtracting negative control values from each treatment group. Error bars represent 95% confidence intervals. All 5 experimental groups were treated with the same dosages of 0, 2.5, 5, 10, and 20 µg/mL WGA. Cell counts for all replicates in ascending order of WGA dosage (see **Table 1** for individual counts): MV3 Det. (470, 602, 479, 501, 562), MV3 Adh. (688, 569, 597, 426, 544), M498 Det. (290, 291, 159, 187, 133), M498 Adh. (405, 378, 347, 285, 316), A375 Det. (867, 797, 670, 847, 679), A375 Adh. (576, 453, 589, 578, 555), Prestressed A375 Det. (496, 493, 649, 317, 325), Prestressed A375 Adh. (495, 581, 448, 504, 475), MAPKi A375 Det. (170,201,240,214,264), MAPKi A375 Adh. (363,374,349,415,354). **(C)** Cell surface renderings of representative cells showing the spatial variation of intracellular mean curvature. **(D)** Cell death upon bleb inhibition using VitroGel coffins for adhered and detached melanoma cells. Cells grown for 24 hours in either integrin-binding VitroGel-RGD (adhered) or non-integrin-binding VitroGel (detached) and assayed for cell death using ethidium homodimer staining. Dots represent individual experiments. Cell counts for all replicates (see **Table 1** for individual counts): MV3 (367, 389), A375 (461, 509).

We performed WGA dose response survival analysis on three different melanoma cell lines: MV3 (NRAS(Q61R)-driven)^34^, A375 (BRAF(V600E)-driven)^35^, and M498 (BRAF(V600E)-driven primary cell line)^36^. After 24 hours of bleb inhibition, detached MV3 cells showed high levels of cell death as measured by ethidium homodimer staining, but little to none when adhered (**Fig 1B**). Both adhered and detached M498 cells die upon WGA treatment, but detached cells experience significantly more death, both in aggregate and in individual matched experiments (**Fig 1B, Fig S1**). Compared to adherent MV3 cells, adherent M498 cells spread poorly and are easily detached with gentle pipetting (**Fig. S2**), suggesting poor adhesion even in growth conditions that permit attachment. Accordingly, bleb inhibition induces an increase in cell death also for the adherent population. In contrast, A375 cells were unaffected by WGA treatment, regardless of their attachment state (**Fig 1B**). During the design of this viability assay we observed that the confluency of A375 cells prior to treatment seemed to influence outcomes, with cells from more confluent culture conditions apparently experiencing more death upon WGA treatment. To examine this systematically, we allowed A375 cells to remain overconfluent for 48 hours (changing culture media daily) before performing viability experiments as before. We found that anoikis resistance in these “pre-stressed” A375 cells was strongly dependent upon blebbing, with an effect size even larger than that seen in MV3 cells (**Fig 1B**). Having established that A375 cells can be made reliant on blebbing by certain stress conditions, we applied the viability assay also to experiments in which A375 cells were challenged by direct pharmacological stress. Specifically, we measured the effect of bleb inhibition on A375 cells pre-incubated for 48 hours in media containing 10nM Dabrafenib (targeting BRAF(V600E)) and 1nM Trametinib (targeting Mek1/2), a combination used as a frontline therapy for Braf-mutated tumors. Drug-treated A375 cells experienced increased cell death upon WGA treatment when detached and minimal change in viability when adhered (**Fig 1B**).

To ensure that the WGA-mediated disruption of anoikis resistance was due to bleb inhibition and not an off-target effect of WGA, we performed a complementary experiment using VitroGel, an abiotic hydrogel used in cell culture^37^, to form bleb-restricting “coffins” around individual cells (**Fig 1C**). Coffins for “detached” cell growth were made using non-adherent VitroGel, while adherent coffins were made using VitroGel-RGD, which contained integrin-binding RGD domains that allow for cell adhesion. We performed these experiments on WGA-sensitive MV3 cells and WGA-insensitive A375 cells. As expected, far more MV3 cells died in non-adherent than in adherent VitroGel coffins, whereas A375 cells showed no difference in viability between the two conditions (**Fig 1D**). Taken together, these results indicate that dynamic blebbing contributes to anoikis resistance both in NRAS- and in BRAF-mutated melanoma, suggesting that this cell morphological program is a broadly adopted survival strategy in melanoma cells. Moreover, the observation of MAPK inhibition (MAPKi) via Dabrafenib/Trametinib sensitizing BRAF(V600E)-driven cells to bleb inhibition suggests that bleb-formation serves also as an acute defense mechanism against drug attacks.

### Bleb-generated plasma membrane curvature drives the formation of cortical septin structures

A notable feature of blebbing vs bleb-inhibited cells is the enrichment of micron-scale plasma membrane curvature. For an intracellular observer, curvature is negative/concave in the bleb proper (reaching values in the range of κ = -1.5 µm^-1^, where κ represents the inverse of the radius of curvature) and positive/convex between the blebs (reaching values in the range of κ = 1.5 µm^-1^) (**Fig 1C**). Serendipitously, we discovered a marked recruitment of septins near densely-packed blebs in MV3 cells expressing murine SEPT6-GFP (**Fig S4**). Indeed, in eukaryotic cells, members of the septin cytoskeletal family are the only proteins known to detect the positive micron-scale membrane curvature that blebbing creates^38,39^. The notion that bleb-generated curvature could recruit septins to the plasma membrane upon cell detachment is supported by previous work ^40,41^, though we found no reports of an explicit connection between bleb-related surface curvature and septin recruitment. Septins are known to scaffold and regulate factors from diverse signaling pathways^42–45^, including pro-survival pathways in cancer cells such as HIF-1^46–48^, EGFR^49^, MET^50,51^, JNK^52^, and HER2^53^. Therefore, we hypothesized that bleb-dependent anoikis resistance could be achieved through recruitment of curvature-sensitive septin proteins to the plasma membrane via bleb-generated curvature, where they promote pro-survival signaling.

To test such an involvement of septins as translators of cell morphology into survival signaling, we performed mass spectrometry of MV3 cell lysate and found robust expression of several septin proteins (**Table 2**). We opted to visualize septin structures in living cells by ectopic expression of mouse SEPT6-GFP^40^ because this construct has been shown to successfully integrate into the endogenous septin cytoskeleton with little-to-no effect on septin function^40,54^. We confirmed proper integration of the fluorescent probe into native septin oligomers by pulldowns of mouse SEPT6-HALO. Mass spectrometry showed that all endogenous septins expressed in MV3 cells were pulled down by SEPT6-HALO, at levels 36x higher than in control pulldowns with no SEPT6-HALO expression (septins accounted for 0.22% of total protein abundance in the control and 8.1% in SEPT6-HALO pulldown), suggesting that the probe efficiently forms oligomers with endogenous septins (**Table 2** and **Supplemental Data 1**). Because previous work has suggested that septins contribute to the retraction of large blebs induced by osmotic shock in T cells, we measured the blebby surface fraction of MV3 cells with and without SEPT6-GFP expression to be sure that our visualization strategy did not itself alter blebbiness. This analysis showed no difference in blebby surface fraction between the two groups (**Fig S5**).

**Table 1.** Cell counts for pertinent experiments.

| Figure 1B | MV3<br>Detached | MV3<br>Adhered | M498<br>Detached | M498<br>Adhered | A375<br>Detached | A375<br>Adhered | Prestressed<br>A375<br>Detached | Prestressed<br>A375<br>Adhered | MAPKi<br>A375<br>Detached | MAPKi<br>A375<br>Adhered |
| --- | --- | --- | --- | --- | --- | --- | --- | --- | --- | --- |
| 0 WGA | 172 | 305 | 121 | 112 | 313 | 137 | 242 | 93 | 24 | 76 |
| 2.5 WGA | 179 | 234 | 107 | 123 | 340 | 162 | 93 | 87 | 20 | 78 |
| 5 WGA | 151 | 254 | 84 | 103 | 234 | 184 | 54 | 75 | 27 | 85 |
| 10 WGA | 109 | 136 | 117 | 94 | 376 | 171 | 31 | 79 | 33 | 101 |
| 20 WGA | 137 | 254 | 69 | 96 | 281 | 181 | 42 | 74 | 28 | 98 |
| 0 WGA | 104 | 167 | 89 | 161 | 212 | 204 | 69 | 59 | 104 | 207 |
| 2.5 WGA | 113 | 185 | 104 | 113 | 134 | 138 | 59 | 51 | 94 | 241 |
| 5 WGA | 123 | 174 | 36 | 137 | 195 | 211 | 99 | 28 | 103 | 200 |
| 10 WGA | 120 | 139 | 38 | 111 | 131 | 162 | 51 | 42 | 108 | 225 |
| 20 WGA | 133 | 149 | 33 | 137 | 153 | 173 | 46 | 57 | 153 | 204 |
| 0 WGA | 194 | 216 | 80 | 132 | 342 | 235 | 185 | 343 | 48 | 80 |
| 2.5 WGA | 310 | 150 | 80 | 142 | 323 | 153 | 341 | 443 | 87 | 55 |
| 5 WGA | 205 | 169 | 39 | 107 | 241 | 194 | 496 | 345 | 110 | 64 |
| 10 WGA | 272 | 151 | 32 | 80 | 340 | 245 | 235 | 383 | 73 | 89 |
| 20 WGA | 292 | 141 | 31 | 83 | 245 | 201 | 237 | 344 | 83 | 52 |

| Figure 1D | MV3<br>Detached | MV3<br>Adhered | A375<br>Detached | A375<br>Adhered |
| --- | --- | --- | --- | --- |
| Rep 1 | 94 | 194 | 152 | 131 |
| Rep 2 | 87 | 114 | 113 | 132 |
| Rep 3 | 91 | 33 | 89 | 108 |
| Rep 4 | 95 | 48 | 107 | 138 |

| Figure 3B | MV3<br>Detached | MV3<br>Adhered | M498<br>Detached | M498<br>Adhered | Unstressed<br>A375<br>Detached | Unstressed<br>A375<br>Adhered | Prestressed<br>A375<br>Detached | Prestressed<br>A375<br>Adhered | MAPKi<br>A375<br>Detached | MAPKi<br>A375<br>Adhered |
| --- | --- | --- | --- | --- | --- | --- | --- | --- | --- | --- |
| Experimental 1 | 170 | 241 | 81 | 111 | 277 | 184 | 345 | 66 | 109 | 245 |
| Control 1 | 194 | 216 | 80 | 132 | 313 | 137 | 230 | 53 | 98 | 364 |
| Experimental 2 | 133 | 149 | 100 | 132 | 205 | 156 | 39 | 70 | 93 | 192 |
| Control 2 | 104 | 167 | 89 | 161 | 212 | 204 | 242 | 93 | 104 | 207 |
| Experimental 3 | 137 | 254 | 117 | 95 | 341 | 222 | 34 | 33 | 29 | 79 |
| Control 3 | 172 | 305 | 121 | 112 | 342 | 235 | 69 | 59 | 31 | 131 |

| Figure 4D | MV3<br>Detached | MV3<br>Adhered |
| --- | --- | --- |
| Experimental 1 | 279 | 256 |
| Control 1 | 453 | 253 |
| Experimental 2 | 388 | 281 |
| Control 2 | 349 | 246 |
| Experimental 3 | 438 | 262 |
| Control 3 | 237 | 235 |

| Figure 4E | Control | FCF |
| --- | --- | --- |
| Adhered | 65 | 61 |
| Adhered | 48 | 59 |
| Adhered | 58 | 53 |
| Adhered | 63 | 63 |
| Detached | 236 | 164 |
| Detached | 212 | 149 |
| Detached | 350 | 258 |
| Detached | 252 | 278 |

| Figure 4F | Adhered | Detached |
| --- | --- | --- |
| Control | 65 | 98 |
| Control | 67 | 73 |
| Control | 75 | 73 |
| Control | 69 | 50 |
| SEPT2(33-306) | 63 | 36 |
| SEPT2(33-306) | 33 | 46 |
| SEPT2(33-306) | 42 | 40 |
| SEPT2(33-306) | 69 | 42 |
| VitroGel | 61 | 84 |
| VitroGel | 56 | 64 |
| VitroGel | 55 | 63 |

| 5B | 90 min | 120 min | 180 min |
| --- | --- | --- | --- |
| MEF | 113 | 67 | 99 |
| MEF+DYN2(K44A) | 42 | 21 | 21 |
| MEF | 67 | 23 | 27 |
| MEF+DYN2(K44A) | 73 | 16 | 70 |

| Figure 5C | Adhered | Detached |
| --- | --- | --- |
| Control | 89 | 49 |
| DYN2(K44A) | 55 | 38 |
| DYN2(K44A)+WGA | 25 | 32 |
| DYN2(K44A)+FCF | 33 | 27 |
| Control | 25 | 23 |
| DYN2(K44A) | 39 | 28 |
| DYN2(K44A)+WGA | 43 | 27 |
| DYN2(K44A)+FCF | 34 | 28 |
| Control | 54 | 92 |
| DYN2(K44A) | 84 | 121 |
| DYN2(K44A)+WGA | 115 | 101 |
| DYN2(K44A)+FCF | 70 | 141 |

**Table 2.** Septin proteomic results.

| Experiment | Septin | % of total septin abundance<br>(fold change between lysate and pulldown) | % of total protein abundance<br>(fold change between lysate and pulldown) |
| --- | --- | --- | --- |
| MV3<br>Lysate | SEPTIN2 | 19.99 | 0.04 |
|  | SEPTIN6 | 40.98 | 0.09 |
|  | SEPTIN7 | 8.80 | 0.02 |
|  | SEPTIN9 | 12.93 | 0.03 |
|  | SEPTIN8 | 0.31 | < 0.01 |
|  | SEPTIN10 | 16.97 | 0.04 |
|  | SEPTIN11 | ND | ND |
|  | SEPTIN3 | ND | ND |
|  | **All Septins** | N/A | **0.22** |
| MV3<br>SEPT6-HALO<br>Pulldown | SEPTIN2 | 30.09 (1.50) | 2.43 (60.75) |
|  | SEPTIN6 | 12.82 (-3.20) | 1.03 (11.44) |
|  | SEPTIN7 | 29.88 (3.39) | 2.41 (120.50) |
|  | SEPTIN9 | 22.97 (1.78) | 1.85 (61.67) |
|  | SEPTIN8 | 0.09 (-3.44) | < 0.01 (N/A) |
|  | SEPTIN10 | 3.19 (-5.32) | 0.28 (7) |
|  | SEPTIN11 | 0.94 (N/A) | 0.08 (N/A) |
|  | SEPTIN3 | ND (N/A) | ND (N/A) |
|  | **All Septins** | N/A | **8.06 (35.86)** |
| A375<br>Lysate | SEPTIN2 | 26.42 | 0.03 |
|  | SEPTIN6 | 31.71 | 0.03 |
|  | SEPTIN7 | 15.60 | 0.02 |
|  | SEPTIN9 | 18.99 | 0.02 |
|  | SEPTIN8 | ND | ND |
|  | SEPTIN10 | 1.92 | < 0.01 |
|  | SEPTIN11 | 5.36 | < 0.01 |
|  | SEPTIN3 | ND | ND |
|  | **All Septins** | N/A | **0.12** |
| A375<br>SEPT6-HALO<br>Pulldown | SEPTIN2 | 25.71 (-1.03) | 1.81 (60.33) |
|  | SEPTIN6 | 23.77 (-1.33) | 1.67 (55.67) |
|  | SEPTIN7 | 21.07 (1.35) | 1.48 (74) |
|  | SEPTIN9 | 22.08 (1.62) | 1.55 (77.5) |
|  | SEPTIN8 | 0.02 (N/A) | < 0.01 (N/A) |
|  | SEPTIN10 | 7.23 (3.77) | 0.51 (N/A) |
|  | SEPTIN11 | 0.12 (-44.67) | < 0.01 (N/A) |
|  | SEPTIN3 | ND (N/A) | ND (N/A) |
|  | **All Septins** | N/A | **7.04 (56.02)** |

Cell surface renderings of SEPT6-GFP indicated that septin structures are primarily found near blebby cell surface regions, while less blebby regions have few-to-none (**Fig 2A, Movie 2**). To confirm this observation quantitatively we applied a directional correlation method^32^, which determines the spatial alignment of cell surface events independent of cell shape. Applied to multiple MV3 cells imaged at isotropic resolution to accurately measure surface curvature and signal intensity in three dimenions^55^, we confirmed association between bleb occurrence and septin localization (**Fig 2B**). Specifically, septins are enriched at positive curvatures above κ=0.4 µm^-1^, in agreement with previous reports of high septin affinity for curvature values κ>0.4 µm^-1 38,39,56^ (**Fig 2C**). Analysis of control and WGA-treated MV3 cells also confirmed that bleb inhibition reduces mean plasma membrane curvature above κ=0.4 µm^-1^ (**Fig 2D**). We further examined the precise localization of septin structures relative to blebs and found them to be enriched at bleb edges, aligned with maximal positive curvature (**Fig 2E**). Septin intensities fall steeply towards the bleb center and more slowly as one moves away from the bleb (**Fig 2E**). To ensure that these bleb-associated septin structures were not an overexpression artifact caused by SEPT6-GFP, we repeated our analyses on MV3 cells that lacked the construct using immunofluorescence. We found that signal from an anti-SEPT2 antibody was enriched at the cell surface near blebs (**Fig S6A-C**), localized to the curvy bases of blebs (**Fig S6D**), and importantly, that SEPT6-GFP expression did not alter anti-SEPT2 enrichment near the surface (**Fig S6E**).

**Figure 2:**
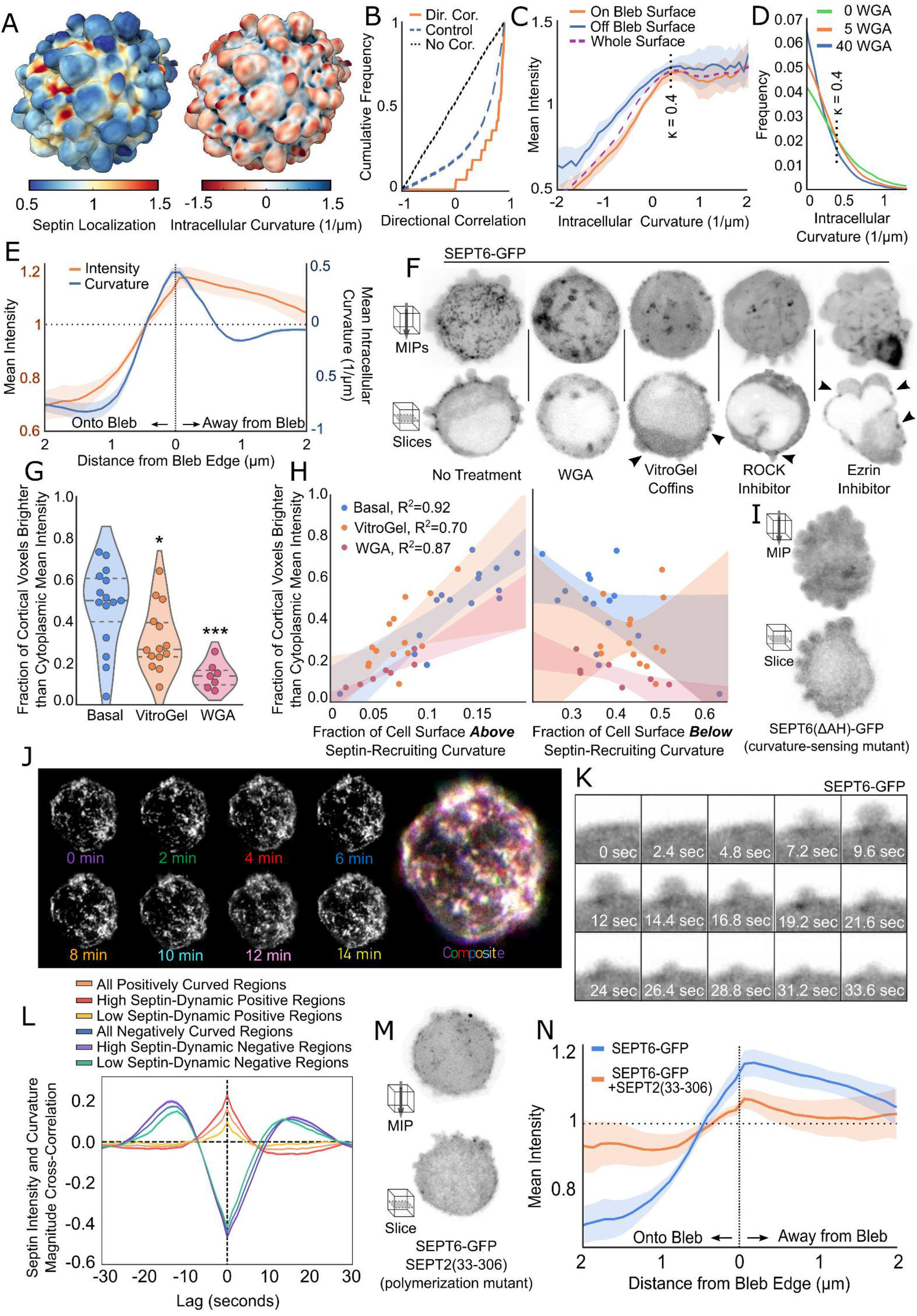
Bleb-generated plasma membrane curvature drives the assembly of cortical septin structures. **(A)** Surface rendering of either the local intensity of the murine SEPT6-GFP probe within 1 µm of the cell surface or local intracellular mean curvature. Cells embedded in soft bovine collagen. **(B)** Directional correlation between blebs and septin localization in 17 MV3 cells and 2025 blebs. Cumulative correlative distribution is shown as orange solid line, randomized bleb localization control is shown as a dashed blue line, and zero correlation is shown as black dotted line. **(C)** Local intensity of septin signal as a function of local intracellular curvature, using same cells. The orange line indicates curvature/intensity relationship on the surface of blebs, the blue line on non-bleb surfaces, and the dashed purple line on the entirety of surfaces. Error bands represent 95% confidence intervals. **(D)** Probability distributions of local positive intracellular mean curvature for all cells in **Fig. 1A** with WGA dosage shown in µg/mL.**(E)** Local septin intensity and intracellular mean curvature as a function of distance from bleb edges using same cells as **Fig. 2B. (F)** Localization of the SEPT6-GFP probe in MV3 melanoma cells with diverse perturbations of bleb formation. Representative cells received treatment as follows: WGA (50 ug/ml), VitroGel, H1152 (1 µM), NSC668394 (10 µM). Maximum intensity projections (MIP) and single z-slices of 0.16 micron thickness. Arrowheads indicate septin accumulation in perturbed cells in regions with residual high-curvature. Cells embedded in soft bovine collagen. **(G)** Fraction of cortical voxels (within 0.96 µm of surface) in basal and bleb-inhibited MV3 cells with septin intensity higher than cytoplasmic mean intensity. WGA group treated at 10 µg/mL. Dashed lines separate quartiles. Basal/VitroGel tested with two sample t-test using pooled variance (p=0.013), normality tested with Shapiro-Wilk (p=0.2579 & 0.31), variance tested with two-tailed F test (p=0.448). Basal/WGA tested with Welch’s t-test (p<0.0001), normality tested with Shapiro-Wilk (p=0.4539 and 0.7941), variance tested with two-tailed F-test (p=0.0114) **(H)** Same data as **Fig. 2G** expressed as a function of cell surface fraction possessing intracellular curvature either above (right, κ > 0.4 µm^-1^) or below (left,0 > κ > 0.4 µm^-1^) septin recruiting threshold (R^2^ p-values = 0.0032, 0.0054, & 0.000027). **(I)** Septin localization in representative MV3 melanoma cell expressing septin curvature-sensing mutant SEPT6(ΔAH)-GFP. **(J)** Time-lapse sequence showing changes in murine SEPT6-GFP probe localization over 14 minutes in maximum intensity projections of a representative MV3 cell. Individual time points shown in grayscale, left; and in a pseudo-color composite, right. Cell embedded in soft bovine collagen. **(K)** Time-lapse showing murine SEPT6-GFP probe accumulation during an individual blebbing event. Single z-slice of 0.16 µm thickness. Cell embedded in soft bovine collagen. **(L)** Overlaid temporal cross-correlation functions between SEPT6-GFP probe intensity and intracellular mean curvature magnitude for contigs comprised of positive (orange/red/yellow) or negative (blue/purple/green) intracellular mean curvature. Colors denote subdivision of datasets depending on dynamicity, as indicated in figure. Data from an MV3 cell embedded in soft bovine collagen imaged for 4 minutes with a stack acquisition rate of ∼0.83 Hz. **(M)** SEPT6-GFP probe localization in representative MV3 melanoma cell expressing septin polymerization mutant SEPT2(33-306). **(N)** In red, local SEPT6-GFP intensity as a function of distance from bleb edges in MV3 cells expressing SEPT2(33-306), overlaid with SEPT6-GFP intensity curve from **Fig 2E** in blue.

To more thoroughly test the notion that bleb formation as a morphodynamic process causally drives the assembly of septin structures, we perturbed blebbing by a variety of methods: In addition to WGA and VitroGel we used the ROCK inhibitor H1152 (reduces bleb formation by lowering intracellular pressure) and the Ezrin-inhibitor NSC668394 (causes large-scale membrane-cortex detachment and thus substitutes single-micron scale, dynamic blebs with large, stable blebs). Every approach had pronounced effects on septin localization, reducing the overall cortical levels with remaining foci confined to high curvature areas (**Fig 2F**). To quantify these observations we measured cortical septin levels as the surface fraction of cortical voxels (voxels <0.96 µm from the surface) above mean cytoplasmic SEPT6-GFP intensity. Both WGA and VitroGel treatments significantly reduced cortical septin levels relative to control conditions (**Fig 2G**). Moreover, all three experimental conditions displayed a strong linear relationship at the single cell level between the surface fraction of predicted septin-recruiting curvature (κ ≥ 0.4 µm^-1^) and cortical septin levels (**Fig 2H**). In contrast, cortical septin levels did not display such a relation to the surface fraction of positive curvature below the septin-recruiting threshold (0 µm^-1^ ≥ κ ≥ 0.4 µm^-1^) (**Fig 2H**). Thus, septin levels at the cortex universally correlate with the presence of septin-recruiting plasma membrane curvature, and experimental conditions which reduce such curvature, like those inhibiting bleb formation, abrogate septin recruitment. To exclude the possibility that septins localize to bleb edges through a process independent of membrane curvature sensing, we generated mutants of our SEPT6-GFP construct lacking the C-terminal amphipathic helix necessary for curvature sensing by septins^38,57^ (**Fig S7**). In contrast to WT SEPT6-GFP, MV3 cells expressing SEPT6(ΔAH)-GFP lacked septin localization to the surface (**Fig 2I, Fig S7**). This demonstrates that the observed cortical septin structures depend entirely on septins’ ability to sense the micron-scale membrane curvature associated with bleb formation.

### Cortical septin structures are formed by repeated bleb-driven curvature events

We found gross septin localization to be relatively stable on a timescale of minutes (**Fig 2J**), despite bleb lifetimes typically falling below 60 seconds^58^. This poses the question of how a relatively brief morphodynamic event can determine longer-lived molecular architectures. To address this, we turned to high-speed time-lapse light-sheet microscopy. These data revealed that while “pulses” of septin accumulation can be seen at the curvy bases of blebs, they fade quickly once the bleb is resolved and curvature subsides **(Fig 2K, Movie 3**). To confirm this observation we computationally identified regions of the surface with high septin intensity that maintained either positive or negative curvature for at least 25 seconds (the average bleb lifetime, estimated by curvature autocorrelation, **Fig S8A**). Within these “contigs” cross-correlation analysis between septin intensity and curvature magnitude showed positive correlation in the positively curved dataset, whereas the negatively curved dataset produced the opposite effect (**Fig 2L, Fig S8B**). Interestingly, when we subdivided contigs based on septin intensity dynamics, we found that more stable regions were less sensitive to positive curvature than dynamic regions, this in contrast to negative contigs, which maintained the same septin/curvature relationship regardless of dynamicity. Examining the mean rates of septin intensity gain or loss within individual contigs as a function of dynamicity showed increased rates of change within more dynamic regions (**Fig S8C**), meaning that the areas in which septin concentration shifts more frequently, those shifts also occur at faster rates. This suggests the presence of two populations within septin localizations at the cell surface: dynamic septin pools dependent upon positive curvature, and more stable pools that are less reliant on specific local curvature profiles.

Cytosolic septins predominantly exist as single hexameric or octameric oligomers that preferentially bind to curved membrane, but quickly detach unless local concentration becomes high enough to promote inter-oligomer polymerization, which stabilizes membrane binding^38,39,59^. If the stable septin regions identified by time-lapse imaging were polymerized higher-order structures, this would predict that not only septin oligomers’ ability to sense curvature, but also their ability to polymerize with one another when membrane-bound above a critical local concentration, is necessary for the formation of bleb-associated septin structures. To test this, we engineered the SEPT2 mutant SEPT2(33-306), which is capable of integrating into septin oligomers but lacks the N- and C-terminal domains necessary for inter-oligomeric polymerization^60,61^ (**Fig S9**). When we observed gross septin localization in MV3 cells expressing SEPT2(33-306), we found that septin enrichment at the surface was strongly inhibited in a dominant negative fashion (**Fig 2M, Fig S9, and Fig S10**). Bleb-adjacent septin localization still showed a sharp peak at the curvy bleb edge, indicating a continuation of curvature-sensing septin pulses, but moving away from the bleb the signal decayed rapidly and to lower levels compared to the signal profiles in unperturbed cells (**Fig 2N**). Conversely, the signal decay was slowed moving towards the center of blebs, matching the decay rate in the opposite direction, and the on-bleb signal was significantly increased. This altered bleb-adjacent localization supports a requirement for inter-oligomer polymerization in the formation of curvature-independent stable septin structures, and suggests polymerized septin structures have an increased affinity for non-bleb plasma membrane. This bleb exclusion might be driven by polymerization-dependent crosslinking of the plasma membrane and actomyosin cortex by septins and associated factors, as has been previously observed^62^.

Indeed, our time-lapse data show that a vast majority of septin pulses endure only as long as the associated blebs, but occasionally a rare de novo formation of stable septin structure can also be observed. In each such instance we noted several pulses occurring in close proximity to one another, resulting in their coalescence into bright long-lived structures. Eventually, their intensities seemed to no longer rely on local curvature profiles (**Movie 4**). Taken together, these findings suggest that stable cortical septin structures are formed by iterative bleb-driven curvature events that result in local levels of septin oligomers surpassing the threshold necessary for inter-oligomer polymerization and enabling stabilization through formation of higher-order structures. This would mean that septins are acting as a discrete time integrator of a sustained and spatially persistent dynamic bleb formation process, while also efficiently eliminating the effects of random and isolated morphological events. Thus, septins are ideally suited to translate significant cell morphological cues into cellular signals.

### Septins are necessary for bleb-dependent anoikis resistance

Given the causal relation between bleb formation and septin localization, we sought to determine whether such cortical septin structures would be implicated in the acquisition of bleb-dependent anoikis resistance. Live cell imaging of poorly adhered cells embedded in soft collagen showed that MV3, M498, pre-stressed A375, and MAPKi-treated A375 cells possess extensive cortical septin structures spatially associated with blebs (**Fig 3A**). In contrast, unperturbed A375 cells, which had shown bleb-independent anoikis resistance (Fig. 1B), displayed no septin enrichment at the cortex despite robust bleb formation.

**Figure 3:**
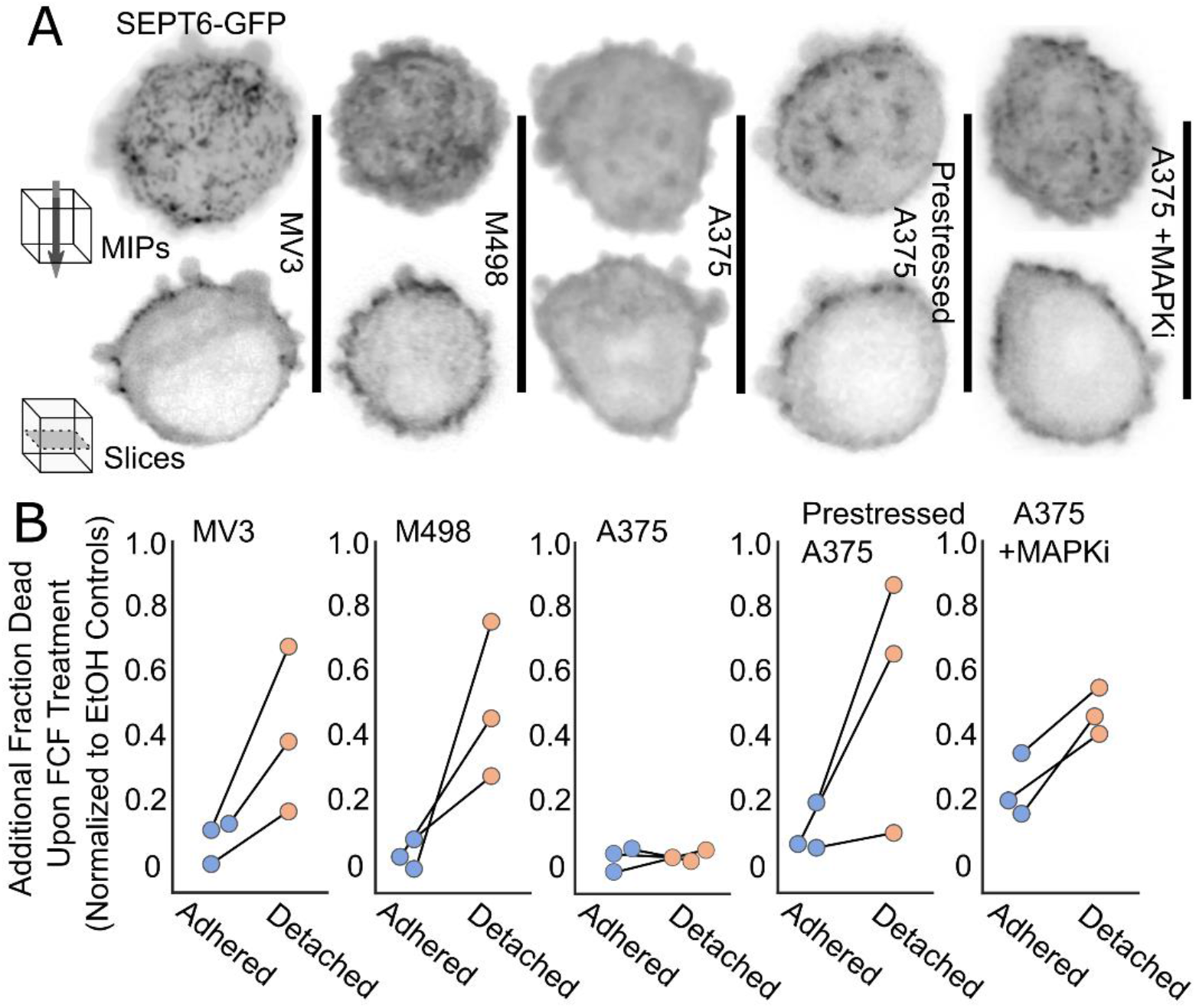
Septins are necessary for bleb-dependent anoikis resistance. **(A)** Murine SEPT6-GFP probe localization in different melanoma cell lines and conditions. Maximum intensity projections (top) and single z-slices of 0.16 micron thickness (bottom) shown for representative cells.. Cells embedded in soft bovine collagen. **(B)** Cell death upon septin inhibition with 50 µM FCF for adhered and detached melanoma cells. Cells grown as in **Fig 1B** and treated with either FCF or EtOH control for 24 hours. Data were normalized by subtracting paired negative control values from each treatment group. Dots represent individual experiments. Cell counts for all, with control counts in parentheses (see **Table 1** for individual counts): MV3 Adh.440(470), MV3 Det 644(688), M498 Adh. 298(290), M498 Det. 338(405), A375 Adh. 823(867), A375 Det. 562(576), Prestressed A375 Adh. 418(541), Prestressed A375 Det. 169(205), MAPKi A375 Adh. 231(233), MAPKi A375 Det. 516(702).

We repeated pull-down mass spectrometry experiments in unperturbed A375 cells and confirmed that the apparent lack of cortical septins was not due to a failure of the probe to integrate into native septin structures (**Table 2**). Moreover, SEPT6-GFP pulldown and whole-lysate mass spec datasets showed no major differences in septin expression patterns or oligomer subunit ratios between MV3 and unperturbed A375 cells (**Table I**). To test for subtle shifts in septin expression between unperturbed and MAPKi-treated A375 cells we performed Western blots measuring changes in the expression of SEPT2, SEPT6, SEPT7, and SEPT9 in MAPKi dose-response experiments. No consistent changes in expression or appearance of alternate bands indicating expression of alternate isoforms could be detected (**Fig S11**). These results suggest that the “septin activation” that occurs upon prestress or MAPKi is not driven by changes in septin expression but is instead post-translationally regulated, likely through control of septin curvature sensing and/or higher-order polymerization. Of note, in neither MV3 nor A375 cells did we find SEPT6 to pull down BORG proteins, a family of septin effector proteins known to regulate higher-order polymerization (**Supplemental Data 2**).

To directly test the conclusion that bleb-associated septin structures regulate survival signaling, we measured anoikis-resistance subject to treatment with the septin inhibitor forchlorfenuron (FCF)^63^. FCF treatment resulted in a near-complete ablation of cortical septin structures without altering blebbiness (**Fig S5**) and, as predicted, greatly disrupted anoikis resistance for MV3, M498, and pre-stressed or MAPKi-challenged A375 cells (**Fig 3B**). In contrast, FCF treatment had no effect on unperturbed A375 cells, mirroring bleb inhibition results. To ensure that these results were not due to off-target effects of FCF we performed an orthogonal experiment using FCF-sensitive MV3 cells and FCF-insensitive A375 cells, inhibiting septin assembly with the dominant negative SEPT2(33-306) mutant. Like FCF, expression of this construct has no effect on blebbiness (**Fig. S5**). As with the FCF treatments, genetic inhibition of septins had no effect on A375 cell survival regardless of adhesion state, while MV3 cells experienced significant death which worsened with detachment (**Fig S12**). Taken together, these data suggest that the mechanism allowing blebs to confer anoikis resistance upon melanoma cells depends on bleb-adjacent cortical septin structures.

### Bleb-associated cortical septins interact with NRAS and MAPK effector proteins

We hypothesized that bleb-induced septin structures promote anoikis resistance as a signaling scaffold that amplifies survival signals. To identify signaling candidates we performed BioID proximity labeling^64^ using SEPT6 as bait. We obtained 321 septin-interacting proteins (full results in **Supplemental Data 1**), among them three plasma membrane-localized components of major signal transduction pathways: Notch, CD44, and NRAS. As Notch and CD44 are canonically juxtacrine, we focused on NRAS as a more likely candidate for survival signaling in detached cells. We also found Prohibitin^65^, 14-3-3ζ^66^, and Nucleolin^67,68^ among the BioID prey, which are all potent effectors of the RAS/RAF/MEK/ERK and RAS/PI3K/AKT pathways. Importantly, reexamination of our previous pulldown data revealed that NRAS interacted with SEPT6 in MV3 and pre-stressed A375, but not in unperturbed A375 (**Supplemental Data 2**), indicating that the NRAS/septin interaction depends on the assembly of bleb-nucleated septin structures. NRAS was also an especially intriguing candidate as MV3 cells harbor a constitutively active Q61R mutant, leaving open the possibility that suspended NRAS-driven melanoma cells harness their mutational profile via an acute morphological program to empower oncogenic signaling through the RAS/RAF/MEK/ERK and RAS/PI3K/AKT axes.

To support these proteomic results, we performed live cell imaging studies to confirm that the SEPT6/NRAS interaction occurs in rounded, detached cells. As expected, ectopically expressed NRAS-GFP strongly co-localizes with cortical septins (**Fig 4A**). Moreover, perturbation of septin structures using FCF or the dominant negative SEPT2(33-306) mutant significantly reduced enrichment of NRAS near the cell surface (**Fig 4B, Fig S13**). To further investigate how septin inhibition affects NRAS localization, we employed Earth Mover’s Distance (EMD)^69,70^ as a metric for the spatial heterogeneity of the NRAS signal at the cell surface, comparing to homogenous distributions of the same amount of signal on the same surface. The results followed our predictions: Septin-inhibited cells produced lower EMD values (suggesting diffuse signal distributions) than unperturbed cells (**Fig 4C**). Taken together, these data strongly suggest that cortical septin structures bind and scaffold NRAS, affecting its spatial distribution near the surface and increasing its cortical concentration.

**Figure 4:**
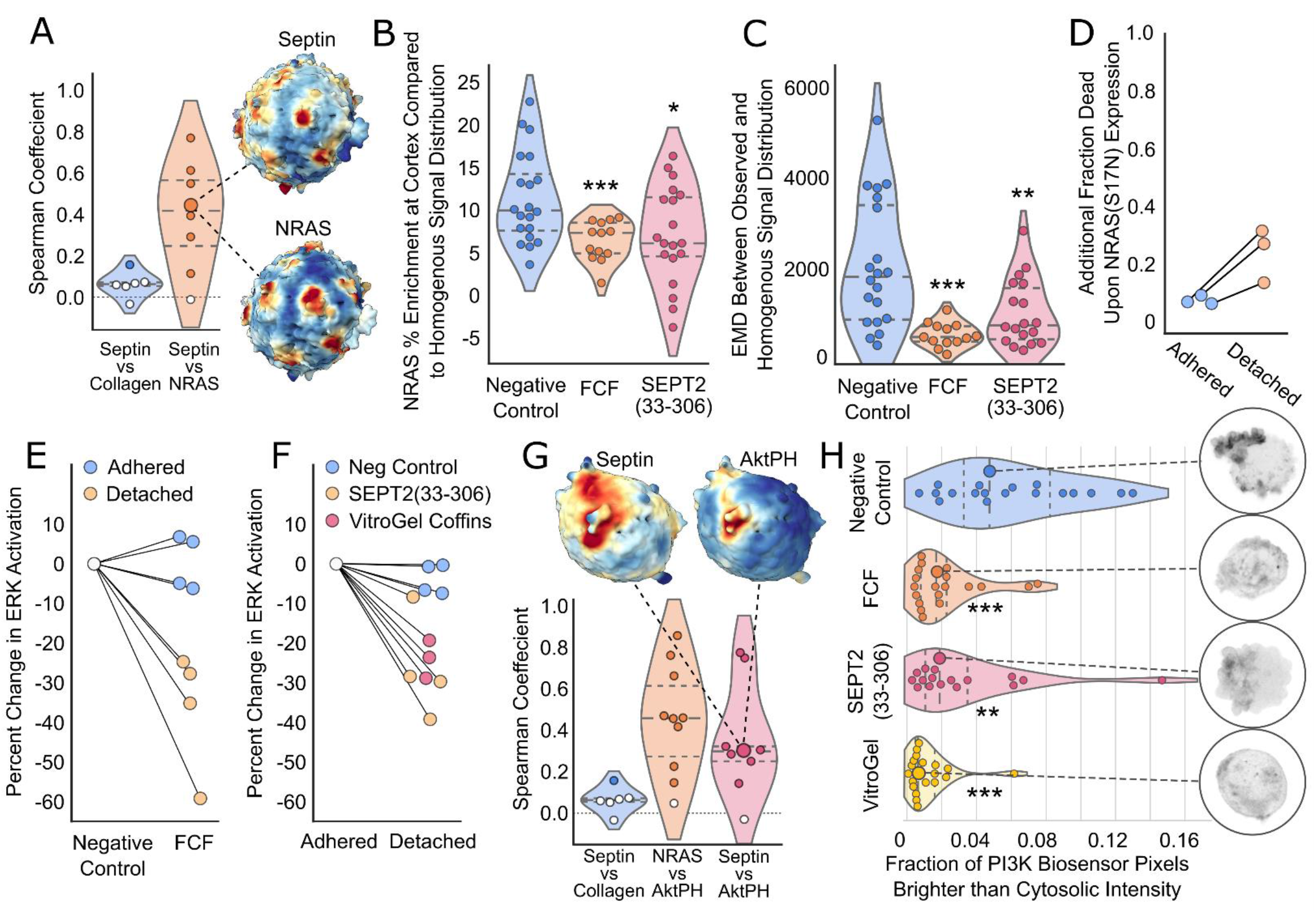
Septins scaffold NRAS, promoting NRAS/MAPK and NRAS/PI3K survival signaling. **(A)** Spearman correlation between NRAS and septin signal distributions on the surfaces of MV3 melanoma cells, compared to septin vs collagen negative control. White datapoints represent cells whose correlations were not significantly higher than analyses performed on the same surfaces with one signal randomly scrambled. Septin and NRAS signal distributions from the median cell in the dataset shown on the right. **(B)** Observed NRAS-GFP % enrichment at the cortex (voxels within 0.96 µm of surface) of individual unperturbed and septin-inhibited MV3 cells. Basal/FCF tested with Welch’s T-test (p<0.0001), normality tested with Shapiro-Wilk (p=0.2191 & 0.2198), variance tested with two-tailed F test (p=0.0054). Basal/SEPT2(33-306) tested with two sample t-test using pooled variance (p=0.0101), normality tested with Shapiro-Wilk (p=0.2191 & 0.7246), variance tested with two-tailed F test (p=0.768).Dashed lines separate quartiles. **(C)** Earth Mover’s Distance (EMD) between observed NRAS-GFP and homogenous distribution of equivalent signal as measured for each unperturbed or septin-inhibited MV3 cell. Control/FCF tested with Mann-Whitney U test (p= 0.000023), normality tested with Shapiro-Wilk (p= 0.07679 and 0.9351). Control/SEPT2(33-306) Mann-Whitney U test (p= 0.0029), normality tested with Shapiro-Wilk (p= 0.07679 and 0.03938). **(D)** Cell death upon expression of dominant negative NRAS(S17N) for adhered and detached MV3 cells. Cells grown as in **Fig 1B** for 24 hours. Data were normalized by subtracting paired negative control values from each treatment group. Dots represent individual experiments. Cell counts for all replicates, with control counts in parentheses (see **Table 1** for individual counts): Adh. 1105(1039), Det. 799(734). **(E)** Effect of septin inhibition upon ERK activation levels in adhered and detached MV3 cells. Percent change in ERK activation between unperturbed and septin inhibited MV3 cells measured by ERK-nKTR-GFP biosensor. Cells grown as in **Fig 1B** for 12 hours. Dots represent individual experiments. Cell counts for all replicates (see **Table 1** for individual counts): Adh. Control 234, Adh. FCF 236, Det. Control 405, Det. FCF 339. **(F)** Effect of detachment upon ERK activation levels in unperturbed, septin-inhibited, and bleb-inhibited MV3 cells. Percent change in ERK activation between attached and detached MV3 cells measured by ERK-nKTR-GFP biosensor. Cells grown as in **Fig 1B** for 12 hours. Dots represent individual experiments. Cell counts for all replicates (see **Table 1** for individual counts): Adh. Control 276, Det. Control 294, Adh. SEPT2(33-306) 274, Det. SEPT2(33-306) 232, Adh. VitroGel 240, Det. VitroGel 205. **(G)** Spearman correlation between NRAS/Akt-PH and Septin/Akt-PH signal distributions on the surfaces of MV3 melanoma cells, compared to septin vs collagen negative control. White datapoints represent cells whose correlations were not significantly higher than analyses performed on the same surfaces with one signal randomly scrambled. Septin and Akt-PH signal distributions from the median cell in the dataset shown above. **(H)** Effect of septin or bleb inhibition upon PI3K activity in individual MV3 cells. PI3K activity measured with the PI3K biosensor Akt-PH-GFP and expressed as fraction of pixels brighter than cytosolic intensity in normalized sum intensity projections. Median cells from all groups shown on the right.

### Septins promote NRAS/ERK and NRAS/PI3K survival signaling

To determine whether NRAS signaling is important for anoikis resistance in MV3 cells, we overexpressed NRAS(S17N), a dominant negative mutant known to disrupt NRAS signaling^71^. Like bleb and septin inhibition, perturbation of NRAS had little effect on the survival of adhered cells, but consistently increased death in detached cells (**Fig 4D**). To determine whether septin organization of NRAS and its effectors affects downstream MAPK signaling, we used the ERK-nKTR nuclear translocation biosensor^72^, which allowed us to monitor the signaling states of individual cells. Because the prevention of cell-cell adhesion requires maintaining ultra-low cell densities, we opted to use quantitative live cell microscopy rather than Western blotting. After 3 hours of septin inhibition with FCF, we found adhered cells to have no difference in ERK signaling compared to controls, while ERK activation was significantly reduced in detached cells (**Fig 4E**). Similarly, we measured the effect of 3 hours of detachment vs. adhesion on unperturbed MV3 cells, MV3 cells expressing the dominant negative SEPT2(33-306) mutant, and MV3 cells grown in Vitrogel coffins. Detachment of uninhibited cells caused little change in ERK activation, while septin and bleb inhibition in detached cells drastically decreased ERK activation (**Fig 4F**). Thus, both septins and blebs are necessary for maintaining ERK signaling in detached, but not in adhered, MV3 melanoma cells.

Finally, because PI3K signaling can be driven by NRAS, and our previous work has shown that bleb inhibition alters PI3K activity^14^, we also tested whether septin-mediated cortical scaffolding of NRAS(Q61R) would drive PI3K signaling in detached MV3 cells – thereby activating a second survival pathway. Using the PI3K activity biosensor Akt-PH-GFP^51^, we found that high PI3K activity co-localizes with both septins and NRAS in rounded, poorly-adhered MV3 cells embedded in soft collagen, in support of septin-scaffolded NRAS signaling through PI3K (**Fig 4G**). We then performed septin inhibition, either through FCF treatment or expression of SEPT2(33-306), and measured the resulting effect upon PI3K signaling. In both cases, PI3K activity fell significantly (**Fig 4H**). Similarly, bleb inhibition with VitroGel coffins greatly depleted PI3K activity, in agreement with our previously published results showing the same effect upon bleb inhibition by WGA^14^ (**Fig 4H**). Altogether, these data show that bleb and septin inhibition specifically reduces both ERK and PI3K activation levels in detached MV3 melanoma cells, supporting the idea that bleb-nucleated cortical septin structures promote anoikis resistance through the scaffolding and upregulation of NRAS/RAF/MEK/ERK and NRAS/PI3K/AKT signaling.

### Prolonging post-detachment blebbing confers bleb- and septin-dependent anoikis resistance in non-malignant fibroblasts

Most metazoan cells will round and begin blebbing upon detachment from substrate (**Movie 1**)^7–13^. Over the course of 1-2 hours this blebbing will subside, as excess membrane is endocytosed away from the cell surface^11,12,29^. Many cancer cells, however, indefinitely sustain blebbing in low- and no-attachment conditions, an amoeboid phenotype seen at the invasive front of tumors *in vivo*^14–19,30^. Having established that this blebbing drives survival signaling in melanoma cells, we wondered whether non-malignant mammalian cells engage in bleb signaling and, specifically, if anoikis resistance can be conferred to non-cancerous cells simply by prolonging post-detachment blebbing.

We first sought to determine whether detached non-malignant cells form bleb-associated septin structures during the 1-2 hour window before bleb attenuation. Examination of mouse embryonic fibroblast (MEF) and human epithelial kidney 293 (HEK) cells expressing SEPT6-GFP revealed bright bleb-associated cortical septin structures at the surface when imaged before bleb attenuation, and an absence of septins at the surface after attenuation (**Figs 5A** and **S14A**). Because attenuation of blebbing in detached cells is primarily driven by dynamin-dependent clathrin-mediated endocytosis (CME)^29^, we analyzed blebbing in MEF cells expressing dominant negative dynamin mutant DYN2(K44A)^73^, expecting that this mild impairment of CME efficiency would prolong post-detachment blebbing. We found that ∼80% of DYN2(K44A)-expressing MEFs maintained blebbiness 3 hours post-detachment, while for control MEFs the blebbing fraction fell below 10% over the same time period (**Fig 5B**).

**Figure 5:**
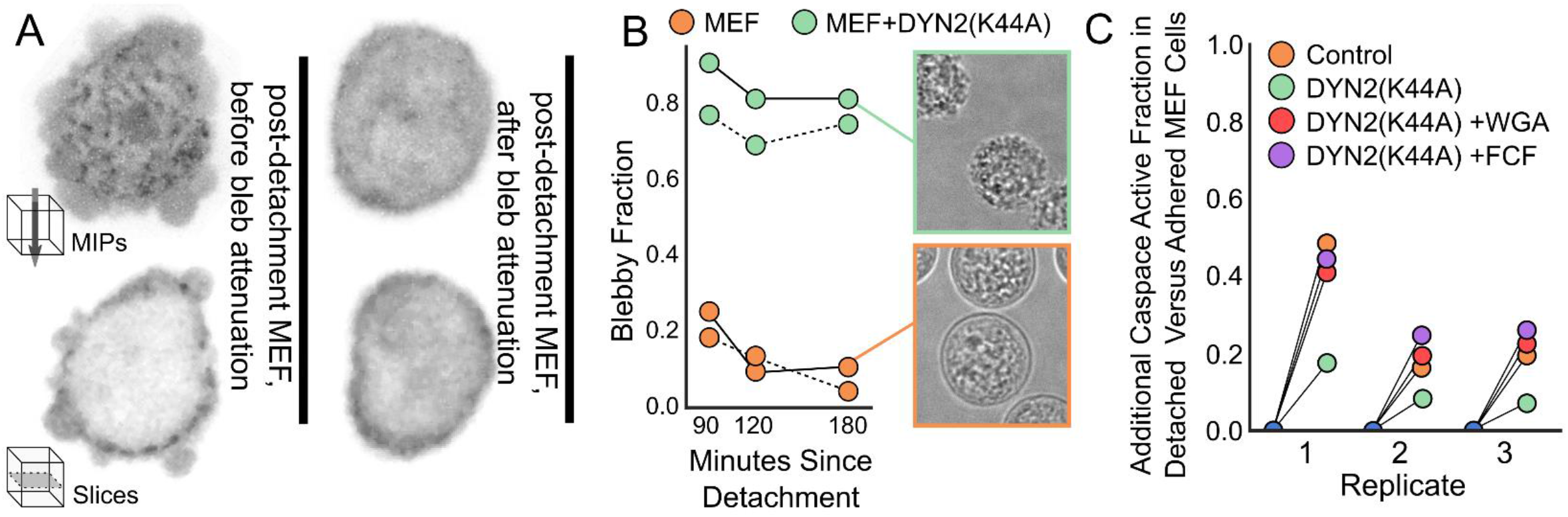
Disrupting bleb attenuation yields anoikis resistance to fibroblasts. **(A)** Recently detached MEF cells expressing SEPT6-GFP, imaged either before or after bleb attenuation. Maximum intensity projections (above) and single z-slices of 0.16 micron thickness (below) shown for representative cells. Cells embedded in soft bovine collagen. **(B)** Fraction of MEF cells showing blebby morphologies 90, 120, and 180 minutes after detachment from substrate. Orange datapoints indicate control MEF cells, green datapoints indicate MEFs expressing DYN2(K44A) in paired, same-day experiments (solid or dashed lines indicate paired data). Representative cells after 180 minutes of detachment shown to the right. Total cell counts for each timepoint, with control group in parenthesis (see **Table 1** for individual counts): 90 min 115(180), 120 min 37(90), 180 min 91(126). **(C)** Additional caspase activation after 4 hours in detached MEF cells compared to paired adhered cells. Orange datapoints indicate control MEF cells, while green datapoints indicate MEFs expressing DYN2(K44A). Red datapoints indicate DYN2(K44A)-expressing cells treated w/ 10 µg/mL WGA and purple indicates DYN2(K44A)-expressing cells treated w/ 50 µM FCF. Caspase activity measured using CellEvent Caspase-3/7 biosensor. Total cell counts for each condition, with adhered group in parenthesis (see **Table 1** for individual counts): Control 164(168), DYN2(K44A) 187(178), DYN2(K44A)+WGA 160(183), DYN2(K44A)+FCF 196(137).

To determine the effects of prolonged blebbiness upon anoikis resistance we measured caspase activity after 4 hours of detachment using the CellEvent biosensor. This indicator of viability was chosen because DYN2(K44A) expression has been observed to reduce cell viability after ≈20 hours of expression (personal communication, M. Mettlen), making it incompatible with the overnight viability assays used in other experiments. To ensure that this assay is able to detect bleb-dependent changes in anoikis resistance, we measured the effect of bleb inhibition on caspase activity in adhered and detached MV3 cells grown for 4 hours. In agreement with our overnight ethidium homodimer viability assays (**Fig 1B**), we found a consistent and specific increase of caspase activation in detached bleb-inhibited MV3 cells (**Fig S14B**).

We then repeated this experiment on MEF cells and found that the blebbing DYN2(K44A)-expressing cells were far more resistant to anoikis than matched same-day controls (**Fig 5C** and **Fig S14C**). To ensure that the difference in caspase activity was not due to an off-target effect of dynamin inhibition, we treated cells with either the bleb inhibitor WGA or the septin inhibitor FCF. In both cases we found that caspase activation was returned to levels seen in matched non-blebbing control cells, meaning that the anoikis resistance acquired by DYN2(K44A) is dependent upon both bleb formation and septins (**Fig 5C** and **Fig S14C**). These data show that cancer cells derived from human melanomas and non-malignant fibroblasts derived from mouse embryos all seem to employ the same bleb and septin dependent anoikis resistance strategy.

## Discussion

Bleb formation has long been recognized as a morphological program associated with metastatic melanoma and other aggressive cancers^14–19,30^. While this program has been largely interpreted as a means of amoeboid migration, our study now shows that dynamic blebbing prompts the construction of septin signaling hubs that substitute for the loss of anchorage-dependent signaling activity upon substrate detachment. Once constructed, such signaling would provide improved survival for cancer cells traveling through low-adhesion environments such as blood, lymph, and stroma. Our experiments with MEFs demonstrate that bleb-based signaling hubs and the anoikis resistance they confer are not oddities specific to melanoma, but rather represent a distinct detachment-activated signaling pathway that is conserved at least across the evolutionary divide between mice and humans, and produces the same pro-survival effect in both normal and malignant cells. While it is unknown what other cell types might employ this bleb/septin signaling paradigm, the widely observed tendency of metazoan cells to bleb upon detachment leads to the conclusion that this strategy might be more broadly distributed and ancient than our current results can unveil.

Septins are highly versatile protein scaffolds that organize and regulate diverse signaling pathways across multiple kingdoms of life^42–45^. While we think our findings support the hypothesis that bleb-associated septins specifically drive anoikis-resistance in melanoma cells via scaffolding of NRAS and downstream pathways, we see no reason to exclude other signaling pathways following the same paradigm. Indeed, according to our proteomic analyses of MV3 cells septins interact with both CD44 and Notch which, despite a canonical involvement in juxtacrine signaling, have both been demonstrated to transduce signal in the absence of activation by neighboring cells or ECM^74,75^. Similarly, dynamic blebbing does not occur only in mammalian cells challenged with detachment, but has also been observed in stem cells^76–79^, migratory cells^14,80,81^, and cells undergoing mitosis^82,83^, not to mention a wide variety of cancer cells^84^. With this in mind, we expect that if future work finds this bleb/septin signaling modality to be important in other cell types, it will not be exclusively RAS-targeting nor survival-oriented, but rather reflect the diversity of cell types and behaviors with which it is associated.

Because MV3 cells harbor the NRAS(Q61R) oncogene classically thought to drive constitutive upregulation of MAPK signaling, our results suggest that this activating mutation is not always sufficient to maintain downstream signaling at levels that guarantee cell viability. Previous work has shown that wildtype and mutant RAS function is regulated through the formation of nanoclusters, small regions of extremely high RAS protein density^85,86^. Our experiments now show that septins serve to modulate local NRAS concentration, thereby affecting downstream signal amplitude. Septin scaffolding is also likely to amplify downstream signals by influencing NRAS interactions with its effectors, such as those found in our proteomic studies. In either case, this finding highlights the functional gap between the genomic condition of possessing an oncogene, and the cellular condition of experiencing penetrant oncogenic signaling. Moreover, it suggests the consideration of septins as a potential therapeutic target for NRAS-mutant melanoma, a disease currently associated with poor prognoses and limited therapeutic options due to the lack of targeted drugs^87^. Forchlorfenuron is a septin inhibitor that has been used as an experimental tool *in vitro* for over a decade^63,88^, with recent studies showing it is effective and well-tolerated in mice^89^. Moreover, because FCF has been in use as a U.S. government-approved agricultural chemical since 2004, EPA studies suggest that it possesses low systemic toxicity in humans^90^, making it (and its chemical derivatives) promising therapeutic candidates.

Our results also demonstrate the potential of such therapeutic strategies against BRAF-mutant melanoma – either as primary therapies as suggested by our M498 results, or adjuvant therapies alongside inhibitors of BRAF(V600E), MEK, and other components of the MAPK pathway as suggested by our A375 results. Our finding of dynamic blebbing conferring resistance to targeted therapies in melanoma agrees with previous work that linked ROCK activation, which directly drives bleb formation by promoting actomyosin contraction, to melanoma therapeutic resistance *in vitro*, in mouse models, and in patients^30^. Furthermore, it was shown that treatment of melanoma cells with MAPKi therapy sensitizes them to blockade of amoeboid morphologies via ROCK inhibition, greatly increasing cell death compared to MAPKi alone. This closely matches our finding that A375 is sensitized to both bleb and septin inhibition by MAPKi, and that co-inhibition of either blebbing or septins produces significantly more cell death than MAPKi alone. Non-genetic adaptive resistance to MAPKi is achieved through the relaxation of negative feedback on NRAS by ERK^91^, with RAF/MEK/ERK signaling remaining low due to sustained MAPKi therapy while survival is maintained by increased NRAS activity feeding into the PI3K pathway. We propose that disruption of bleb-formation and/or cortical septin assembly, like in MV3 cells harboring constitutively active oncogenic NRAS, renders insufficient the signal produced by the less basally active WT NRAS in MAPKi-resistant BRAF(V600E) cells. Support for this proposal is provided by our SEPT6 pulldown experiments, which show NRAS/septin interactions in “septin positive” MV3 and pre-stressed A375 cells, but not in “septin negative” basal A375 cells that lack such structures. The nature of this “septin activation” in A375 cells remains mysterious. Our data shows no changes in septin expression between conditions, indicating post-translational regulation of septins at play. Future work will be necessary to understand the mechanism driving this disease-relevant regulation of the septin cytoskeleton.

In the bleb/septin/NRAS signaling pathway described here, information flow is transduced from chemical (morphogenic signaling), to spatial (blebbiness), and back to chemical (survival signaling), with septins acting not only as a spatiochemical translator, but also as discrete time integrator of pulsatile spatial information. While control of morphology is often thought of as a terminal objective of cell signaling, consideration of cellular morphodynamics as an upstream input to signal transduction defines a new paradigm. Such morphological cues would not only act as potent drivers of signaling, but also as integrators of cell-autonomous and cell external information flows. For example, dynamic blebbing is typified by low Rac1 activity, high Rho/Rock activity, low ERM engagement, high actomyosin contraction, low glycocalyx density, and low attachment^81,92^. All this information is combined into a state that is interpretable as a single signal transduction node by septins, which can then be activated to form versatile signaling hubs capable of regulating myriad signaling pathways.

## Methods

### Cell Culture and Reagents

MV3 cells were a gift from Peter Friedl (MD Anderson Cancer Center, Houston TX). A375 cells (ATCC® CRL1619) were acquired from ATCC. M498 cells were acquired from Sean Morrison (UT Southwestern, Dallas, TX). Mouse embryonic fibroblasts (MEF/3T3) cells were acquired from BD Biosciences, Clontech (# C3018-1). MV3, A375, M498, and MEF cells (were cultured in DMEM (Gibco) supplemented with 10% fetal bovine serum (FBS; ThermoFisher) at 37° C and 5% CO2. We performed targeted deep sequencing of 1385 cancer-related genes showing sequence and copy number variations in MV3, A375, and M498, the results of which are publicly available^93^.“Pre-stressed A375” cells were generated by allowing A375 cells to grow to confluency and then grown for an additional 48 hours without passaging, with media changed daily. “MAPKi A375” cells were generated by treating A375 cells with a combination of 10 nM Dabrafenib and 1 nm Trametinib (alternate concentrations used where noted in the text) for 48 hours.

### Inhibitors

Wheat Germ Agglutinin (WGA) was purchased from Sigma (product # L9640). Forchlorfenuron (FCF) was purchased from Sigma (product # C2791). VitroGel and VitroGel-RGD were purchased from TheWell Bioscience (sku #s TWG001 and TWG002). H1152 was purchased from Tocris (catalogue # 2414). NSC6683394 was purchased from Sigma (product # 341216). Dabrafenib (GSK2118436), and Trametinib (GSK1120212) were purchased from Selleckchem. Transient expression of dominant negative Dyn2(K44A)^94^ was achieved through adenovirus transduction^95^ as previously described^96^ with cells analyzed 16-18 hours after transduction.

### Recombinant DNA Constructs

Mouse SEPT6-GFP construct was purchased from Addgene (Addgene plasmid# 38296) and was cloned into the pLVX-IRES-puro vector (Clontech). The GFP-AktPH construct was obtained from the laboratory of Jason Haugh (North Carolina State University, Raleigh NC) (Haugh et al., 2000) and cloned into the pLVX-IRES-puro vector (Clontech). The GFP-tractin construct was a gift from Dyche Mullins (Addgene plasmid # 58473) and was cloned into the pLVX-IRES-puro vector (Clontech). The BioID2 construct was obtained from Addgene (Addgene plasmid # 74223) and cloned onto the N-terminus of SEPT6 from SEPT6-GFP-pLVX-IRES-puro, replacing eGFP but maintaining the same 22 amino acid linker. SEPT6-HALO was made by cloning the HALO tag from pFN21K (Promega cat# G2821) onto the N-terminus of SEPT6 from SEPT6-GFP-pLVX-IRES-puro, replacing eGFP but maintaining the same 22 amino acid linker. ERK-nKTR-GFP was purchased from Addgene (Addgene plasmid # 59150). C-terminally mGFP-tagged human NRAS in pLenti-C-mGFP was purchased from OriGene (OriGene cat# RC202681L2). C-terminal mGFP was removed and eGFP tag was cloned onto the N-terminus after aberrant localization was observed upon expression in MV3 cells (presumably due to steric inhibition of C-terminal palmitoylation and farnesylation domains by the C-terminal mGFP tag). NRAS(S17N) mutant was generated by cloning the S17N mutation into untagged NRAS-pLenti construct using HiFi assembly. pBOB-Septin2-GFP was purchased from Addgene (Addgene plasmid # 118734), aa 1-32 and 307-361 were removed via PCR to create SEPT2(33-306), and the construct was cloned into pLVX-IRES-puro vector (Clontech). SEPT6(ΔAH)-GFP was generated by removing aa 355-372 of SEPT6 from SEPT6-GFP-pLVX-IRES-puro via PCR. H2B-mCherry was obtained from Addgene (Addgene #89766). Cells expressing lentiviral vectors were created by following the manufacturer’s instructions for virus preparation and cell infection (Clontech). Cells were selected for expression by treatment with puromycin, G418, or by using fluorescence activated cell sorting.

### Detached/Adhered Cell Culture

Cells were grown to 70-80% confluency, trypsinized for 3 minutes, resuspended at approximately 100 cells/ml (measured by eye using light microscopy) in DMEM (Gibco) supplemented with 10% fetal bovine serum (FBS; ThermoFisher) and detached from one another by repeated pipetting. Drug treatments, solvent-only controls, or CellEvent caspase activity sensor were added to suspensions if appropriate to the experiment, and then immediately transferred to both an uncoated (for detached cells) and ibiTreated (for adhered cells) 8-well ibidi µ-slide (ibidi cat# 80821 and 80826) at 200 µl/well. Both slides were stored at 37° C and 5% CO2 for either 24hrs for viability assays, 3 hrs for ERK activity assays, or 4 hours for caspase activation assays. Uncoated slides (detached) were nutated at 20 rpm during this time. At the end of this period detached cells were examined by microscopy to confirm that cells did not aggregate – if aggregates were found the experiment was discarded and repeated. Results from protocol optimization show this was most commonly caused by wells being seeded at too high a cell density. For experiments using MAPKi A375 cells, Dabrafenib and Trametinib treatment was continued over the course of the experiment.

### Cells Embedded in 3D Collagen

Collagen gels were created by mixing bovine collagen I (Advanced Biomatrix 5005 and 5026) with concentrated phosphate buffered saline (PBS) and water for a final concentration of 2 mg/mL collagen. This collagen solution was then brought to pH 7 with 1N NaOH and mixed with cells just prior to incubation at 37° C to induce collagen polymerization. Cells were suspended using trypsin/EDTA (Gibco), centrifuged to remove media, and then mixed with collagen just prior to incubation at 37° C to initiate collagen polymerization. To image collagen fibers, a small amount of collagen was conjugated directly to AlexaFluor 568 dye and mixed with the collagen sample just prior to polymerization. Poorly-adhered cells were enriched in samples by visualizing immediately upon collagen polymerization, and visually selected for those displaying rounded, blebby morphologies.

### Cells Grown in VitroGel Coffins

VitroGel coffins were prepared by embedding MV3 or A375 cells in VitroGel 3D or RGD at 1:1 dilution according to manufacturer’s instructions (TheWell Biosciences, sku # TWG001). Briefly, cells were grown to 70-80% confluency, trypsinized, and diluted to approximately 500 cells/ml in in DMEM (Gibco) supplemented with 10% fetal bovine serum (FBS; ThermoFisher) and detached from one another by repeated pipetting. A 1:1 solution of VitroGel and VitroGel Dilution Solution was made, and added to cell suspension at a ratio of 4:1 (VitroGel:cells) using gentle pipetting until well-mixed (with care taken not to form bubbles). Cell/VitroGel solution was transferred to an 8-well uncoated ibidi µ-slide (ibidi cat# 80821) at 100 µl/well, spread over the bottom of wells with a pipette tip, and allowed to polymerize at room temperature for 15 minutes. The remainder of the well was then filled with 10% FBS DMEM and the slide was stored at 37° C and 5% CO2 for either 24 hrs for viability assays, 3 hrs for ERK activity assays, or 1 hr for visualization of septins / PI3K activity.

### 2D Immunofluorescence

WT MV3 or MV3 cells expressing SEPT6-GFP were seeded in MatTek glass bottom coverslip dishes (P35G-1.5-14-C) at 50,000 cells and incubated overnight. The following day the cells were washed 3 times with 1X PBS and fixed with 37oC pre-warmed 4% PFA dissolved in cytoskeletal buffer. The cytoskeletal buffer consists 10mM PIPES, 100mM NaCl, 300mM Sucrose, 1mM EGTA, 1mM MgCl2, 1mM DTT and Protease inhibitor cocktail. The cells were incubated with PFA at 37oC for 20 mins. The MV3-Sept6-GFP cells were then imaged on the Nikon Eclipse Ti widefield epifluorescence microscope with 100X Plan APO oil immersion lens. For immunofluorescence staining of wild type unlabeled cells, the cells were then permeabilized with 0.5% Triton X-100 for 20 mins and then blocked with 5% BSA in 1X PBS for 1 hour. Following blocking the cells were then treated with rabbit-anti-SEPT2 primary antibody (Sigma, HPA018481) at a 1:1000 dilution overnight at 4oC. The cells were then washed with 1X PBS for 10 minutes for a total of three times and then treated with donkey anti-rabbit-488 secondary antibody (Thermo, A-21206) at a final dilution of 1:5000 for 1 hour. The cells were then washed with 1X PBS for 10 minutes for a total of three times and then imaged on the Nikon Eclipse Ti widefield epifluorescence microscope.

### 3D immunofluorescence

MV3 cells were harvested from 10cm dishes and seeded on Corning Ultra-Low Attachment culture dishes (Sigma, CLS3262) for ∼2 hours for the cells to acquire their blebby architecture. The cells were then centrifuged at 1000 rpm for 5 minutes. The pellet was gently washed once by re-suspending in 1X PBS to remove any media and centrifuged. The pellet was then re-suspended and fixed in 100 µL of pre-warmed 37oC, 4% PFA dissolved in cytoskeletal buffer for 20 minutes at 37oC. The cytoskeletal buffer consists 10mM PIPES, 100mM NaCl, 300mM Sucrose, 1mM EGTA, 1mM MgCl2, 1mM DTT and Protease inhibitor cocktail. The cells were then washed with PBS, and centrifuged. To permeabilize the cells pellet was resuspended in saponin for 20 minutes. The cells were then centrifuged and the pellet was blocked by re-suspending in 5% BSA for one hour with gentle rocking for uniform blocking. The cells suspension was then centrifuged and re-suspended in rabbit anti-SEPT2 primary antibody (Millipore Sigma, HPA018481) at a concentration of 1:1000 overnight at 4°C. The following day the cells were centrifuged, and washed once in 10 mL of PBS. The cells were then centrifuged and re-suspended in donkey anti-rabbit-488 secondary antibody (Thermo, A-21206) at a final dilution of 1:5000 for 1 hour. The cells were then centrifuged and washed twice with 10mL 1X PBS by centrifugation. The pellet was then re-suspended in AF568-phalloidin (Invitrogen, A12380) for one hour, washed, DAPI for 5 minutes, washed, and embedded in soft bovine collagen as described above.

### Viability, ERK Activity, and Caspase Activity Cell Counting Assays

Viability, ERK activity, and caspase activity assays were analyzed using live-cell fluorescence and phase-contrast microscopy, performed on a Nikon Ti microscope equipped with an environmental chamber held at 37° C and 5% CO2 at 20x magnification. For viability assays, cells were stained with ethidium homodimer-1 (Invitrogen cat # E1169) at 4 µM and Hoechst (ThermoFisher cat # H3570) at 10 µg/ml for 15 minutes before imaging. Live and dead cells (as identified by significant cellular ethidium signal) were counted using the Cell Counter ImageJ plugin. For ERK activity assays, live cells carrying ERK-nKTR-GFP and H2B-mCherry were imaged after 3 hours of experimental conditions and high / low ERK activity cells were counted with Cell Counter ImageJ plugin. Cells in which nuclear GFP signals were by eye higher than cytoplasmic signal (“brighter” nucleus visible within “darker” cytoplasm) were labelled ERK low, while cytoplasmic signals equal to or higher than nuclear signal (nucleus indistinguishable from cytoplasm or “darker” nucleus within “brighter” cytoplasm) were labelled ERK high. To control for unconscious bias in making this determination, the identity of all ERK experimental groups were blinded from the analyst. For caspase activity assays, cells were treated with CellEvent caspase-3/7 green detection reagent (ThermoFisher cat # C10423) according to manufacturer’s instructions at a final concentration of 8 µM. Cells were treated upon introduction to adhered or attached culture conditions, as described above. After 4 hours, cells were treated with Hoechst (ThermoFisher cat # H3570) at 10 µg/ml for 15 minutes and imaged. Caspase positive and negative cells (as identified by significant cellular CellEvent signal) were counted using the Cell Counter ImageJ plugin.

### Proteomics

MV3, MV3 expressing SEPT6-HALO, A375, and A375 expressing SEPT-Halo cells were each plated in 150 mm dishes at approximately 7.5 × 10^6^ cells per dish and grown to 70-80% confluency. Pulldowns were performed per manufacturer’s instructions (Promega, G6504). Cells were washed and harvested in ice cold PBS. Cells were pelleted at 2000 RCF for 10 minutes at 4°C. Pellets were stored at -80°C for 72 hours. Pellets were then thawed at room temperature, lysed in Mammalian Lysis Buffer supplemented with Protease Inhibitor cocktail (Promega). The lysate was homogenized with a 27G syringe and centrifuged at 14,000 RCF for 5 min at 4°C. The supernatant was diluted in TBS and incubated with pre-equilibrated HaloLink Resin (Promega) at room temperature with rotation for 15 minutes. Resin was then washed 4 times with Resin Wash Buffer (Promega). Complexed proteins were eluted in SDS Elution Buffer (Promega) for 30 minutes at room temperature. Eluted samples and whole cell lysate controls were loaded and run on a 10% Mini-PROTEAN TGX protein gel (Biorad), visualized with AcquaStain (Bulldog Bio), excised, and analyzed with an Orbitrap Fusion Lumos using reverse-phase LC-MS/MS. MS data were analyzed using Proteome Discoverer 2.2 and searched using the human protein database from Uniprot.

For BioID proximity labeling, approximately 4 × 10^7^ cells were incubated in DMEM (Gibco) supplemented with 10% fetal bovine serum (FBS; ThermoFisher) and 50 µm Biotin for 16 hours at 37° C and 5% CO2. Cells were washed twice in 1x PBS and lysed with 1:1 dilution of 2xJS buffer (100mM HEPES ph7.5, 300mM NaCl, 10mM EGTA, 3mM MgCl2, 2% glycerol, 2% triton-100) containing HALT phosphatase-protease cocktail (ThermoFisher #23225). Cells were collected using a cell scraper, Triton X-100 was added to 2%, and the resulting mixture was put on ice and sonicated. An equal volume of chilled lysis buffer was added and the mixture was sonicated again before centrifugation at 16,500 RCF for 10 minutes. The supernatant was collected and incubated overnight with Dynabeads (ThermoFisher #65602) at 4° C. Beads were magnetically collected and supernatant was removed. Beads were washed 4x with 50 nM Tris-Cl, pH 7.4 with 8 M Urea and supernatant was removed completely. Beads were resuspended in Laemmli buffer and biotinylated proteins were eluted by boiling for 5 minutes. Supernatant was loaded and run on a 10% Mini-PROTEAN TGX protein gel (Biorad), visualized with AcquaStain (Bulldog Bio), excised, and analyzed with an Orbitrap Fusion Lumos using reverse-phase LC-MS/MS. MS data were analyzed using Proteome Discoverer 2.2 and searched using the human protein database from Uniprot.

### Western blots

Cells were lysed in RIPA buffer (50 mM Tris HCl, 150 mM NaCl, 0.5% (w/v) Sodium Deoxycholate, 1.0 mM EDTA, 0.1% (w/v) SDS, 1.0% (v/v) NP-40, and 0.01% (w/v) sodium azide; pH of 7.4). Proteins were run on precast gels (BIO-RAD #4568123) and transferred onto PVDF membranes. After transfer, the membranes were rinsed in TBS buffer, air dried at room temperature for 20-30 minutes, and rewet in TBS buffer supplemented with 0.5% Tween-20 (TBST). The membranes were then blocked for 30min in 5% bovine serum albumin (BSA) for septins or in 5% nonfat dry milk for vinculin. The blocking BSA or milk were dissolved in TBST. Membranes were incubated rocking in blocking solutions with primary antibodies overnight at 4°C, while the the secondary antibodies were rocked at room temperature for one hour. We used the following primary antibodies: SEPT2 (Sigma #HPA018481, 1:1000), SEPT6 (Sigma #HPA005665, 1:1000), SEPT7 (Sigma #HPA023309, 1:1000),

SEPT9 (Sigma #HPA042564, 1:1000), and Vinculin (Santa Cruz Biotechnology #sc-25336, 1:1000). The secondary antibodies used were goat anti-mouse (Invitrogen #31430, 1:20,000) and goat anti-rabbit (Invitrogen #G21234, 1:20,000). Proteins were detected using enhanced chemiluminescence reagents (Thermo Scientific #34095) and imaged on the PVDF membrane (Thermo Scientific #88518) using the G:Box imager (SYNGENE) and GeneSys software. Protein band densitometry was performed in ImageJ.

### 3D Light-Sheet Imaging

3D samples were imaged using two variants of axially-swept light-sheet microscopy^97,98^, the first of which provides sub-400 nm isotropic raw resolution, and the second near-isotropic at ∼400×400×450 nm, uniformly maintained throughout large fields of view of ∼100×100×100 microns. The first variant is equipped with 40X NA 0.8 Nikon illumination and detection objectives, and the second is equipped with a NA 0.67 Special Optics illumination objective and a 25X NA 1.1 Nikon detection objective. For very fast imaging, where aberration-free remote focusing of the illumination light-sheet becomes rate-limiting (faster than approximately 0.1 Hz full volume acquisition, depending on cell size), these microscopes could also be operated in traditional light-sheet microscopy mode. Here, the numerical aperture of the illumination beam was reduced to cover a field of view of ∼20 microns, and imaging was performed by scanning the illumination light-sheet synchronously in the Z-direction with the piezo mounted detection objective.

Samples were imaged in phenol red free DMEM containing 25mM HEPES (ThermoFisher) with 10% FBS and antibiotic-antimycotic (Gibco), held at 37° C during imaging. Images were collected using sCMOS cameras (Orca Flash4.0 v2, Hamamatsu) and the microscopes were operated using custom Labview software. All software was developed using a 64-bit version of LabView 2016 equipped with the LabView Run-Time Engine, Vision Development Module, Vision Run-Time Module and all appropriate device drivers, including NI-RIO Drivers (National Instruments). The software communicated with the camera via the DCAM-API for the Active Silicon Firebird frame-grabber and delivered a series of deterministic TTL triggers with a field programmable gate array (PCIe 7852R, National Instruments). These triggers included analog outputs for control of mirror galvanometers, piezoelectric actuators, laser modulation and blanking, camera fire and external trigger. All images were saved in the OME-TIFF format. The microscope control software is freely available to academic and nonprofit institutions upon completion of a material transfer agreement with the University of Texas Southwestern Medical Center.

### 3D Cell Image Analysis

Cell morphology and septin localization were analyzed principally via u-shape3D^32^, with all exceptions (figures 2G, 2H, 2L, 4A, 4B, 4G, 4H, S5A, and S5B) described below. Briefly, 3D images were first deconvolved using either a Richardson-Lucy or Wiener algorithm with an experimentally measured point spread function. Next, cells were segmented from the image background using u-shape3D’s ‘twoLevel’ mode, which combines a straightforward Otsu threshold^99^ of the image to detect the outer cell surface with a blurred version of the image to segment the inside of the cell. The volume segmented from the blurred image was morphologically eroded to ensure the fidelity of the overall segmentation. Cell surfaces were then represented as triangle meshes, and the mean surface curvature at every triangle was calculated as previously described^32,100^. To remove irregularities, the curvature is next smoothed in real space with a median filter of 1 pixel to remove infinities and then slightly diffused along the mesh.

Septin and NRAS localization was measured from fluorescence images by extending a sphere of either 1 µm (for septins) or 2 µm (for NRAS) about each mesh triangle and mapping to the triangle the mean intensity of intracellular signal within that sphere. Unlike in previous studies^32,100^ the image intensity was not depth normalized prior to analysis, but was instead measured from the raw, undeconvolved image.

Blebs were also detected as previously described^32^. Machine learning models trained via images labeled by three separate expert annotators were combined via voting to classify blebs. Distances from bleb edges were then calculated as the geodesic distance from each triangle to the nearest bleb edge. The blebby surface fraction was also defined as the percentage of total mesh triangles classified as ‘on bleb’. Bleb and septin directional correlations were calculated using spherical statistics^32^, in particular by fitting spherical normal distributions to distributions defined at each mesh triangle.

Cortical septin levels were quantified by measuring the mean cytoplasmic intensity of each cell using hand-drawn ROIs that excluded nuclei (performed with the FIJI ImageJ package), and then calculating for each cell its fraction of cortical voxels (intracellular voxels within 0.96 µm of the u-Shape3D-derived surface) that were higher than that cell’s mean cytoplasmic intensity (performed using a MATLAB script). NRAS enrichment at the surface was quantified by first calculating the total amount of intracellular signal expected to be within 0.96 µm of the u-Shape3D-derived surface if the cell’s signal was homogenously distributed between all voxels, and then calculating the percent change between this value and the observed value for each cell (performed using a MATLAB script). To quantify PI3K biosensor fluorescence signal, cells were segmented with u-shape3D and the intracellular signal was summed across the z-axis to yield a sum projection image (performed with the FIJI ImageJ package). To account for differential biosensor expression levels, projections were normalized by adjusting brightness until the mean cytoplasmic signals of all images were approximately the same. PI3K activity was then quantified as the fraction of total pixels with intensities brighter than a threshold value (the approximate upper range of cytoplasmic signal), which was held constant across cells (performed with the FIJI ImageJ package).

Colocalization of fluorescent signal distributions across 3D cell surfaces was quantified by calculating the Spearman’s rank correlation coefficients between signals on a cell-by-cell basis. To approximate the significance of these correlations a variation of Costes’ randomization^101,102^was employed, in which one distribution was randomized and the Spearman coefficient was calculated again. This process was repeated 1000 times, noting the final fraction of randomized Spearman coefficients greater than that calculated for the data as observed. This fraction is referred to as a *P*-value (not to be confused with the p-value output by statistical tests of significance). Because the sampling of the local intensity of a surface signal through overlapping spheres (as done with u-Shape3D) produces a smoothing effect that leads to spatial autocorrelation, signal distributions were downsampled to below the level of spatial correlation to assure independence of data points. This was accomplished by selecting 200 random equidistant points on a cell’s surface^103^ before Costes’ randomization was performed on these points. To assure that this random selection did not bias the analysis, the process was repeated 10 times, with the resulting *P*-values averaged to a single mean *P*-value. If the mean *P*-value was less than 0.05 the colocalization of the tested signals was deemed significant.

To quantify NRAS distributions, Earth Mover’s Distance (EMD) was measured on discrete surfaces^70^. For a single cell, the measured intensity of NRAS-GFP signal at the surface was modified in two steps to compensate for cell-to-cell variation in fluorescence intensity. Because we were interested only in bright areas representing high NRAS density, a surface background defined as the median of surface intensity was subtracted from the measured surface intensity, and the resulting values were normalized to the mean cytoplasmic intensity derived manually as described above. Resulting negative intensity values along the triangle mesh surface were set to zero. For each cell, the EMD measured the distance between the modified NRAS signal distribution on each cell surface and homogenous distribution of the same amount of signal on the same surface, i.e. the larger the EMD the more spatially organized is the tested signal. These operations were performed using a MATLAB script.

### Analysis of surface curvature and septin timeseries

The goal of these timeseries analyses was to test the central hypotheses that septin intensity co-fluctuates with curvature and that septin intensity increases on stable structures or associated with de novo assembly are significantly enriched on positive intracellular curvature. Substantial pre-processing was required to bring volumetric timelapse datasets of SEPT6-GFP fluorescence into a format enabling such analyses. The processing steps fell into two categories, as detailed below: i) conversion of raw data into analytically tractable mesh datasets (including surface segmentation, registration, curvature measurement, intensity measurement, bleach correction, and smoothing); and ii) identifying portions of these datasets suitable for statistical inference of the coupling between curvature and septin accumulation (including gating curvature, intensity, and intensity change data, and isolating contiguous regions of the surface that exhibit the desired characteristics for significant amounts of time). Once prepared, we assessed the coupling of septin intensity and curvature fluctuations by cross-correlation on contiguous regions of negative and positive intracellular curvature, measured signal dynamicity within these regions, and determined the relationship between septin signal dynamicity and the magnitude of septin intensity changes within these regions.

#### Registration

To extract timeseries, the segmented mesh from the first frame (3D vertex coordinates, ***x***_**0**_ = (*x*_0_, *y*_0_, *z*_0_) and face connectivity) must be consistently tracked over time. To achieve this, we first removed whole-cell motion with rigid-body registration. We then applied non-rigid diffeomorphic registration^104^ on the pre-registered video to infer the individual voxel-wise geometrical (Δ*x*_*t*_, Δ*y*_*t*_, Δ*z*_*t*_) translation vector of all individual frames at time *t* relative to the first frame. The vertex coordinates of the tracked mesh at each individual timepoint *t* are then given by vector summation,(*x*_*t*_, *y*_*t*_, *z*_*t*_) = (*x*_0_, *y*_0_, *z*_0_) + (Δ*x*_*t*_(***x***_**0**_), Δ*y*_*t*_(***x***_**0**_), Δ*z*_*t*_(***x***_**0**_)) relative to the vertex coordinates of the first frame mesh.

#### Curvature and Intensity Measurement

The continuous mean curvature at each vertex position is estimated by quadric fitting^105^ of the vertex coordinates in a 5-vertex ring neighborhood (∼1 µm radius). The corresponding septin intensity was calculated by extending a trajectory to an absolute depth of 1 µm along the steepest gradient of the distance transform to the mesh surface, and assigning the 95^th^ percentile of intensity sampled along that trajectory to the originating vertex to capture the systematically brightest accumulation of septin signal in the cortical shell.

#### Bleach Correction

The raw septin intensity suffers decay from bleaching. We simultaneously normalized and corrected the intensity by computing the normalized septin intensity as the raw intensity divided by the mean septin intensity in the whole cell volume at each timepoint. This normalized septin intensity was used for all subsequent analyses. The instantaneous change in the normalized septin intensity (Δ*I*_sept_) was computed by finite differences between consecutive timepoints at corresponding vertex positions.

#### Smoothing

The raw extracted timeseries are stochastic. Smoothing is required to identify temporally continuous regions of negative and positive curvature from the timeseries. Individual vertex timeseries were temporally smoothed using a moving average with the window size, *w*_*autocorr*_, set by the lag of inflection point in the mean temporal autocorrelation curve (**Fig S7A**). For Δ*I*_sept_, a smoothed timeseries was defined by linear regression of all values *I*_sept_ (*t)* in the closed time interval 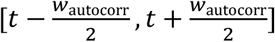. All timeseries were then smoothed spatially using Laplacian smoothing with the number of smoothing iterations inferred by the lag position of the inflection point in the decay of the mean spatial autocorrelation function, which in our data was approximately 1 µm, or a 5-ring neighborhood. Finally, the vertex timeseries were converted to mesh face timeseries using barycentric interpolation.

#### Curvature Gating

Using the mean value of individual timeseries, we derived unbiased thresholds for determining if a face had a positive, flat or negative curvature using 3-class Otsu thresholding.

#### Intensity Gating

Similar to the curvature gating, we applied 3-class Otsu thresholding to the mean intensity value of all smoothed mesh face septin timeseries *I*_sept_, to identify depleted, background, and enriched septin intensity values. For Δ*I*_sept_, 3-class Otsu thresholding was applied to the data between 0-10% and 90-100% of each timeseries, respectively, to identify timepoints of decreasing and increasing intensity. 3-class thresholding was used to identify nonsignificant, unsure and significant decreasing/increasing intensities.

#### Identifying Contiguous Surface Regions (Contigs) Exhibiting Significant Intensity Change

The above thresholds enable us to functionally annotate the mesh face timeseries at every timepoint and identify only the subset of faces with significantly fluctuating septin. We do this by counting the number of significant septin intensity increases and decreases and applying binary Otsu thresholding. For this face subset, we extract temporally continuous periods of positive and negative curvature using the criteria established above. These periods are referred to as *contigs*. Only contigs greater than *w*_*autocorr*_ were then used to compute the normalized cross-correlation curve between the raw septin and curvature timeseries in the contig.

#### Timeseries Analysis

The mean cross-correlation curve and 95% confidence interval for contigs of positive and negative intracellular curvature were computed to test the extent of co-fluctuation between septin and curvature on faces of negative and positive curvature (**Fig 2L and S7B**). We further bipartitioned the contigs of positive and negative intracellular curvature by a score of dynamicity, which describes the total number of septin increase and decrease events divided by the duration of the contig (*T*_contig_), i.e.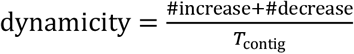. Binary Otsu thresholding was applied to distinguish contigs with low versus high dynamicity. To test how dynamicity relates to Δ*I*_sept_ on negative and positive curvature faces, we computed the continuous relationship between the mean absolute increasing and decreasing Δ*I*_Sept_ vs dynamicity (**Fig S7C**) in contigs using kernel density analysis. Gaussian kernel density with a bandwidth set by Scott’s rule was used to derive the joint density distribution of Δ*I*_Sept_ and dynamicity, i.e. *p*(*X, Y)*, with *X*: dynamicity, *Y*: Δ*I*_Sept_ over the closed intervals *X* ∈ [ 0, 1 ] and *Y* ∈ [ 0, 0.1 ]. The continuous relationship is then given by the marginal expectation, with capital letters denoting the random variable and 𝔼[•] the expectation operator, 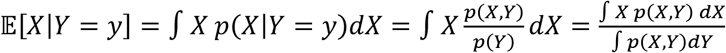 with standard deviation equivalently defined as the square root of the variance,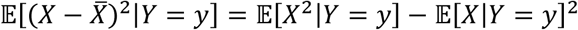. The evaluation of the integrals uses 100 bins for both dynamicity and Δ*I*_sept_.

### Visualization and Statistics

3D surfaces were rendered by ChimeraX^106^. Colored triangle meshes representing the cell surface were imported into ChimeraX from u-shape3D as Collada dae files, as previously described^32^. The FIJI ImageJ package (https://imagej.net/Fiji) was used to prepare MIP and slice images. High-speed 3D light-sheet imaging movies were prepared using Arivis4D. Gardner-Altman paired mean difference estimation plots were generated using DABEST software suite^107^. All other figures were prepared using the Seaborn^108^ and Matplotlib^109^ Python visualization libraries, and the pandas^110,111^ Python data analysis library. Figures were assembled using the InkScape (https://inkscape.org/) vector graphics editor. Significance tests were performed using either two sample t-tests with pooled variance, Welch’s T-tests, or Mann-Whitney *U* tests, depending on whether datasets had normal distributions (as measured by Shapiro-Wilk tests) and equal variance (as measured by F-tests). Statistical calculations were performed using R for Windows, 4.0.5 (https://www.r-project.org/). Error bars in figures show 95% confidence intervals. Number of cells and/or number of different experiments analyzed are given in the figure legends and in Table 1.

## Data Availability

The data that support the findings of this study are available from the corresponding author upon reasonable request.

## Code Availability

All code/algorithms/software central to the findings of this study are available from the corresponding author upon reasonable request. Most can be found at https://github.com/DanuserLab.

## Acknowledgements

We would like to thank John Huang of TheWell Bioscience for providing information about the physical properties of VitroGel, Dagan Segal for discussions about septin inhibition, Xuexia Jiang for conversations about optimal transport, Philippe Roudot for conversations on 3D image analysis, Tadamoto Isogai for advice on proximity proteomics, Benjamin Nanes for assistance in data blinding, and Michael McMurray for helpful feedback on the manuscript. Funding for this work in the Danuser lab has been provided through grants R35 GM136428 (NIH) and I-1840-20200401 (Welch Foundation). Andrew Weems is a fellow of the Jane Coffin Childs Memorial Fund.

---

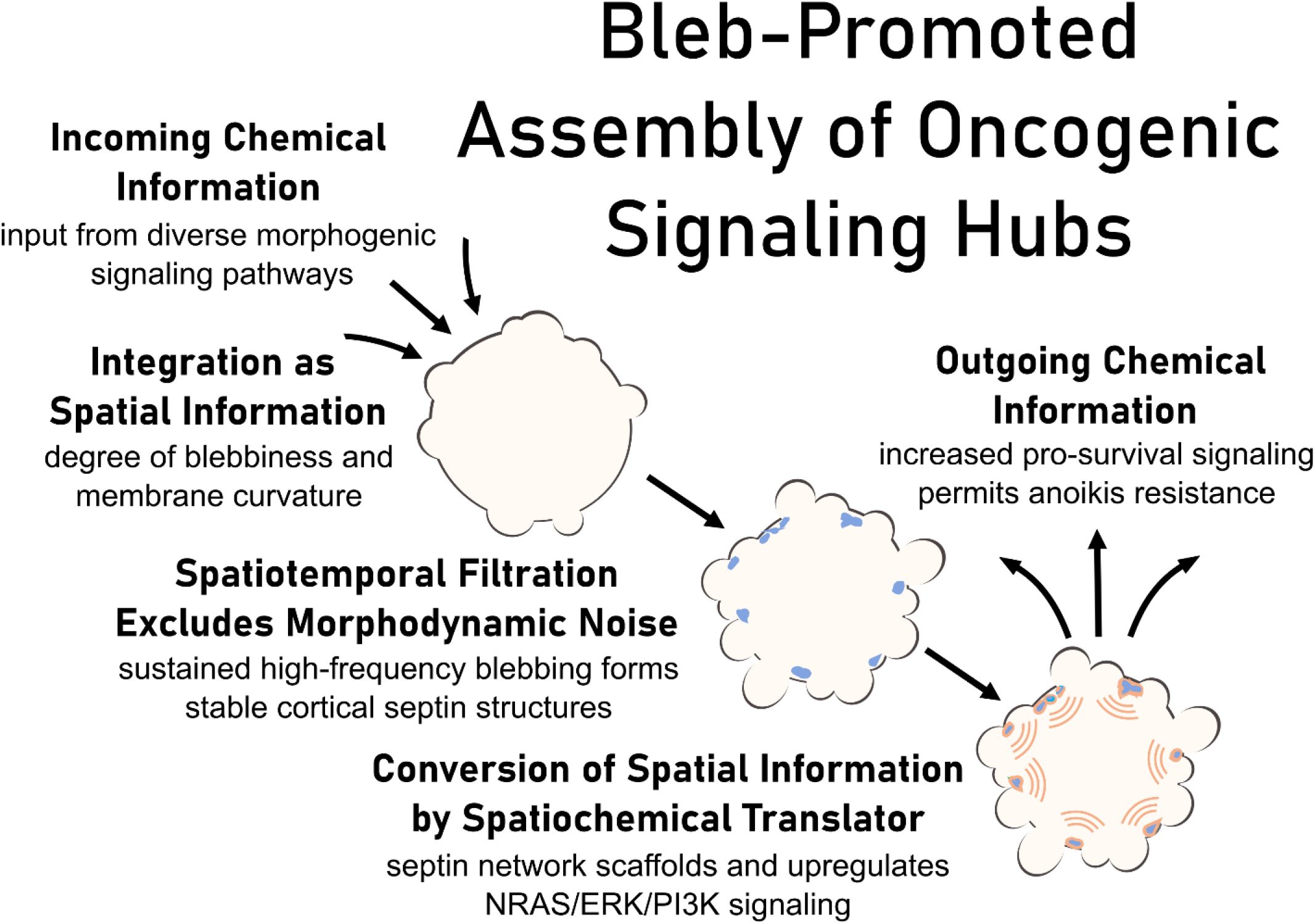

**Figure S1:**
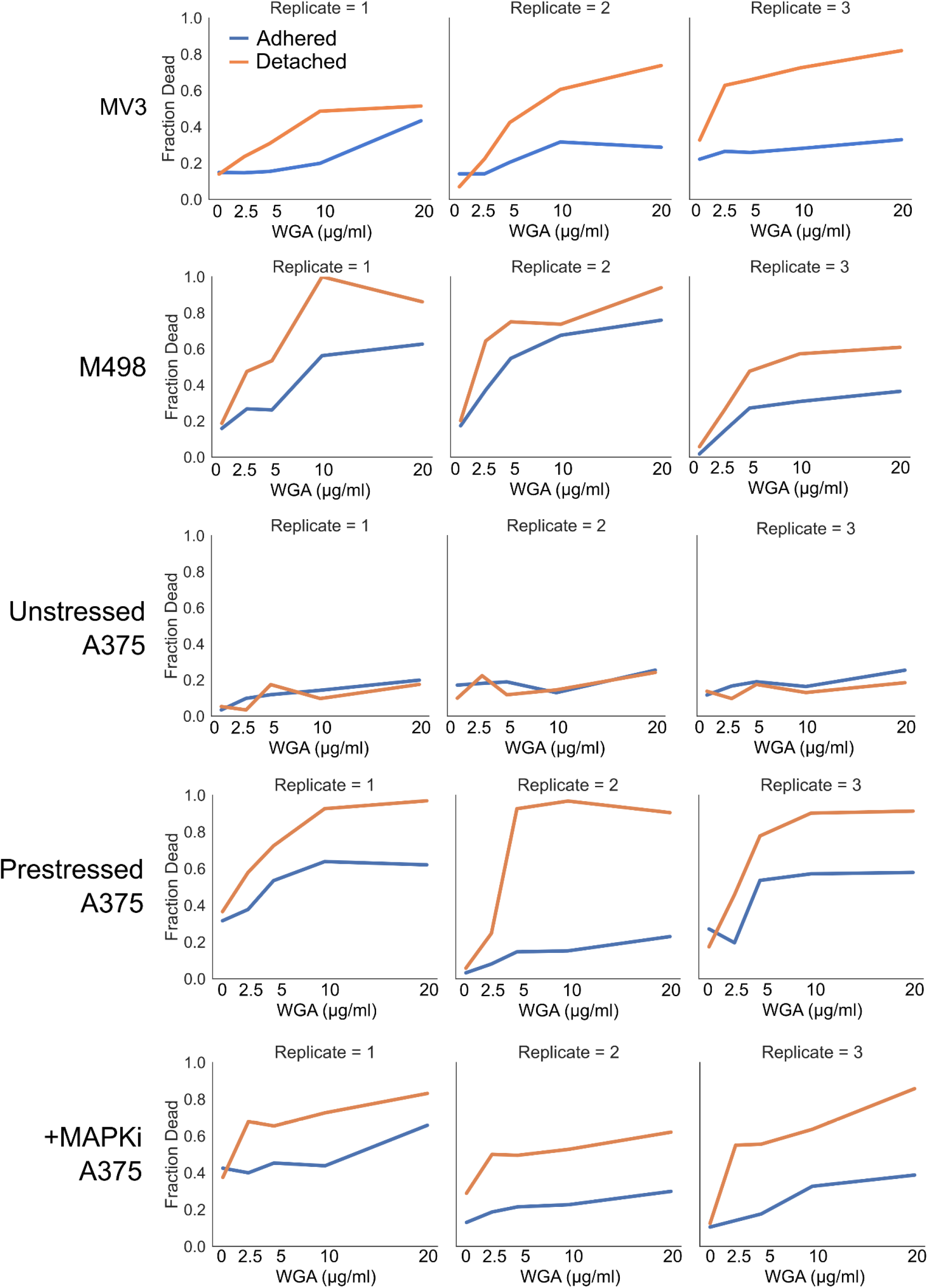
Replicates of bleb inhibition viability experiments. Individual bleb inhibition viability experiments, as summarized in **Figure 1B**. Each pairing of adhered/detached cell experiments was performed on the same day and seeded with cells from the same diluted cell suspensions.

**Figure S2:**
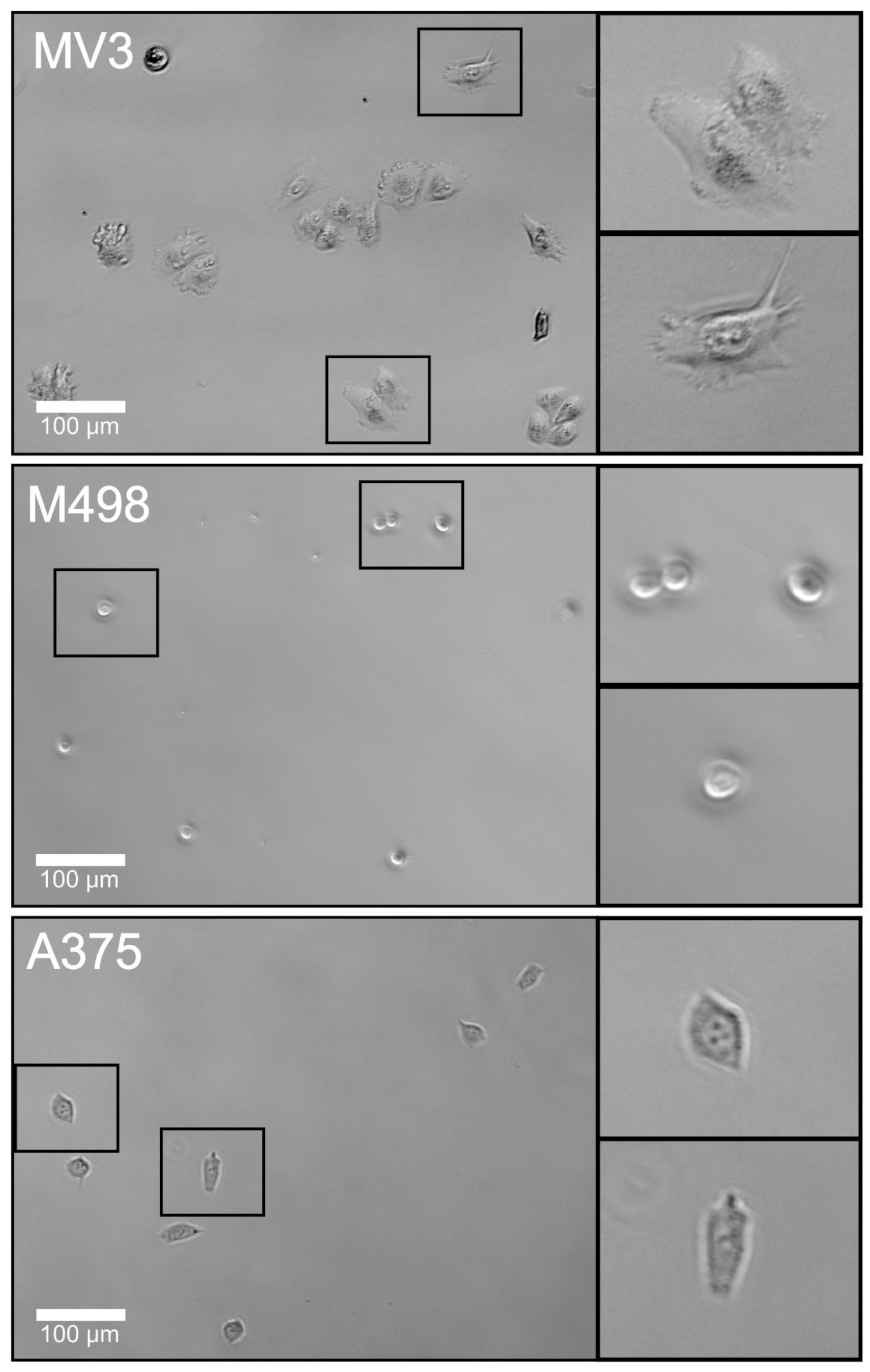
Morphology of cells in “adhered” conditions. Cells grown without perturbation for 24 hours on adherent chamber slides.

**Figure S3:**
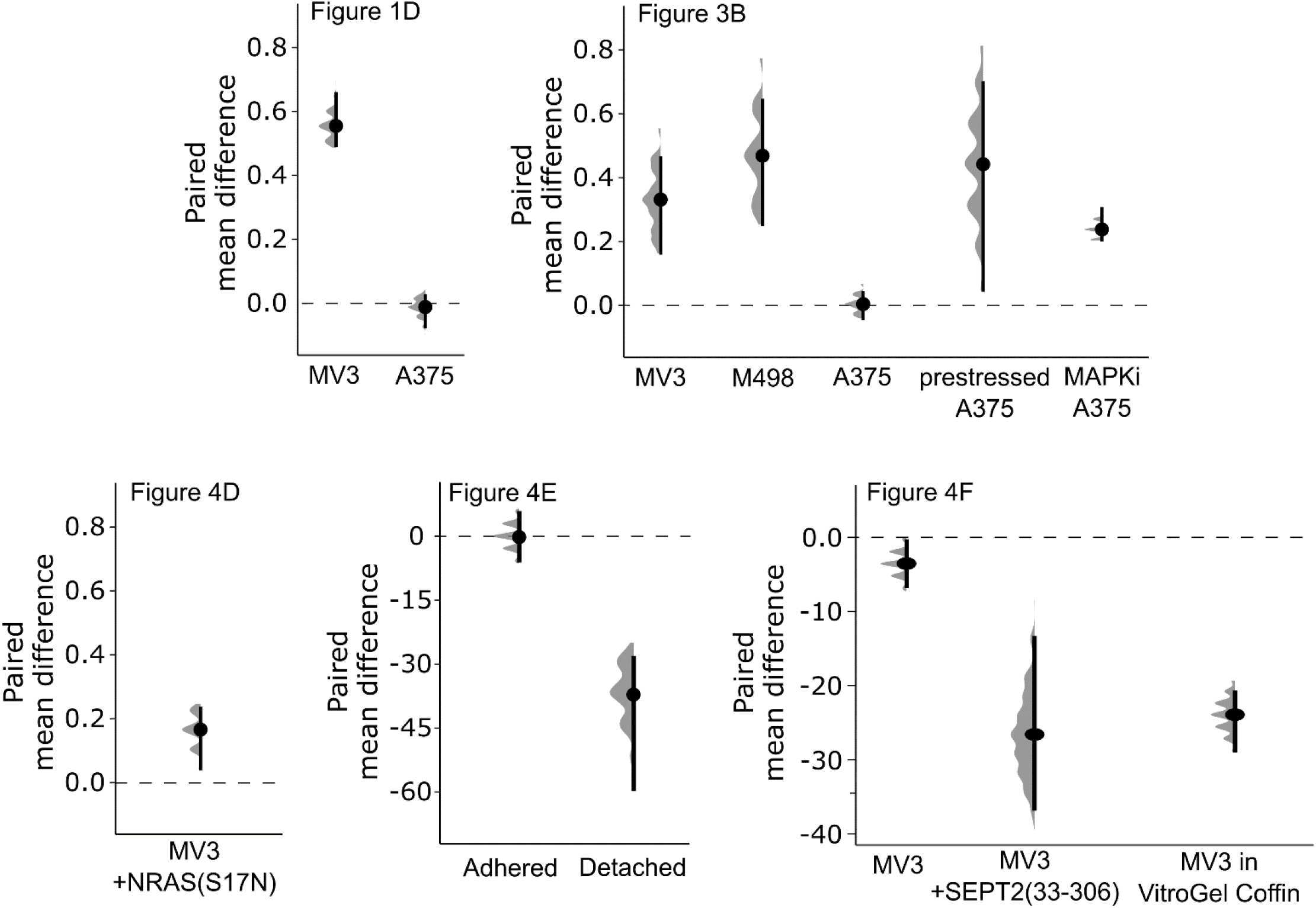
Paired mean differences between control and experimental condition. shown in Gardner-Altman estimation plots for the data presented in figures **1D, 3B, 4D, 4E**, and **4F**. Gray distributions represent 5000 bootstrapped samples, black bars represent 95% confidence intervals, black dots represent mean difference.

**Figure S4:**
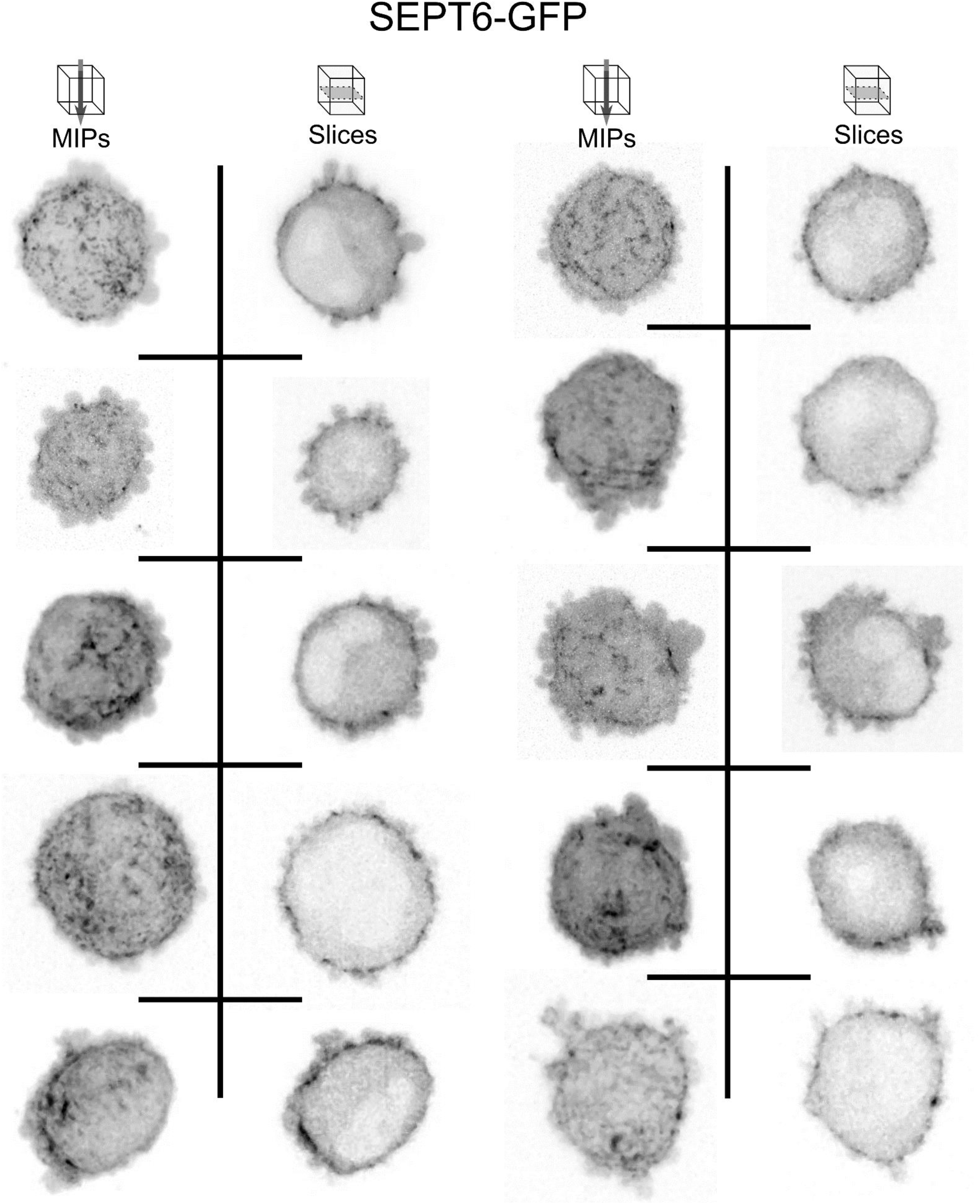
SEPT6-GFP localization in MV3 cells. Murine SEPT6-GFP probe localization in MV3 cells embedded in soft bovine collagen. Maximum intensity projections (left) and single z-slices of 0.16 micron thickness (right) shown for representative cells in two columns.

**Figure S5:**
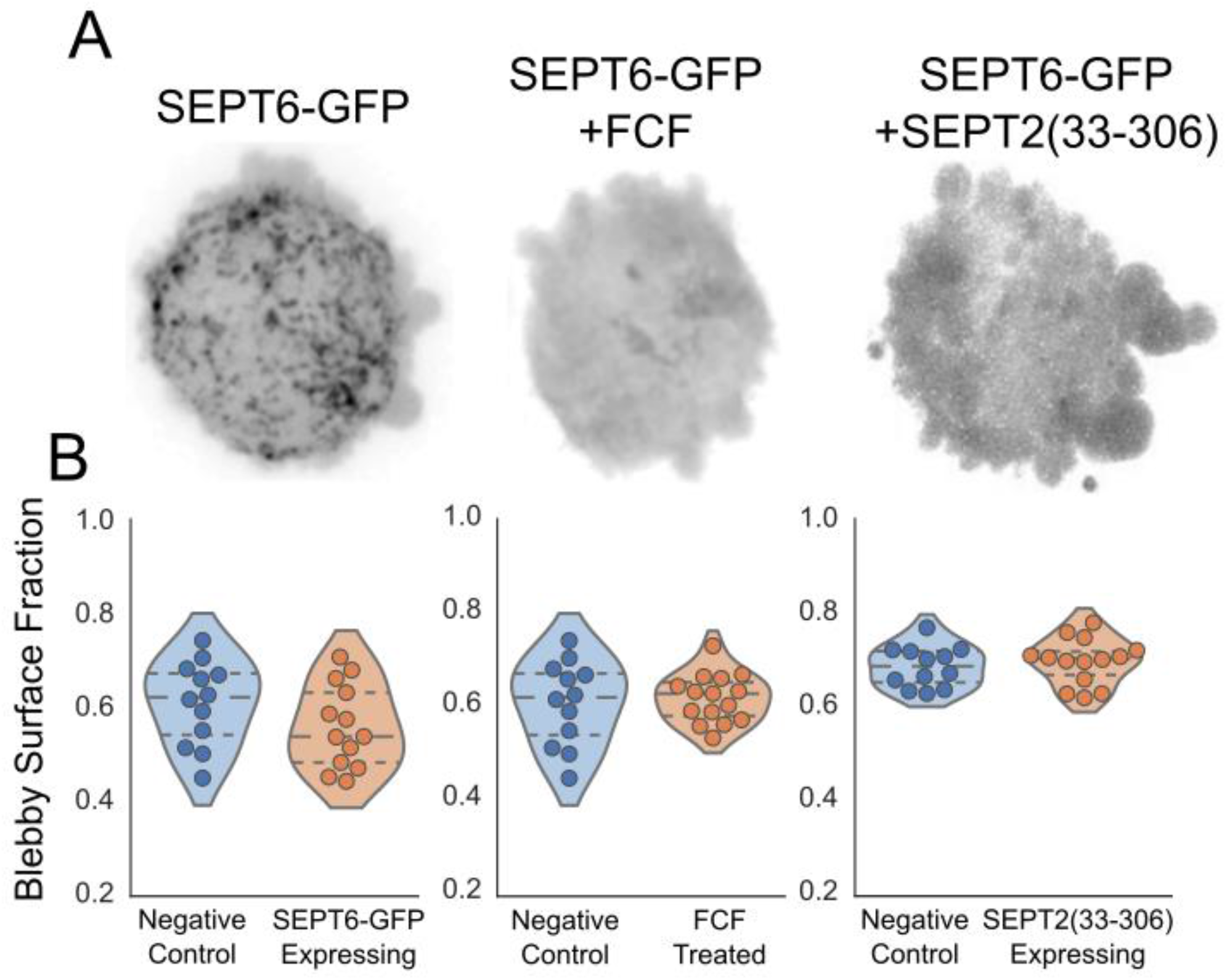
Effects of SEPT6-GFP expression and septin inhibition upon cell blebbiness. **(A)** Maximum intensity projections of murine SEPT6-GFP probe localization in MV3 cells with and without inhibition by FCF or SEPT2(33-306) expression. Cells embedded in soft bovine collagen. **(B)** Fraction of cell surface comprised of blebs for MV3 cells with and without SEPT6-GFP expression, FCF treatment, and SEPT2(33-306) expression. Dashed lines separate quartiles. Dots represent individual cells.

**Figure S6:**
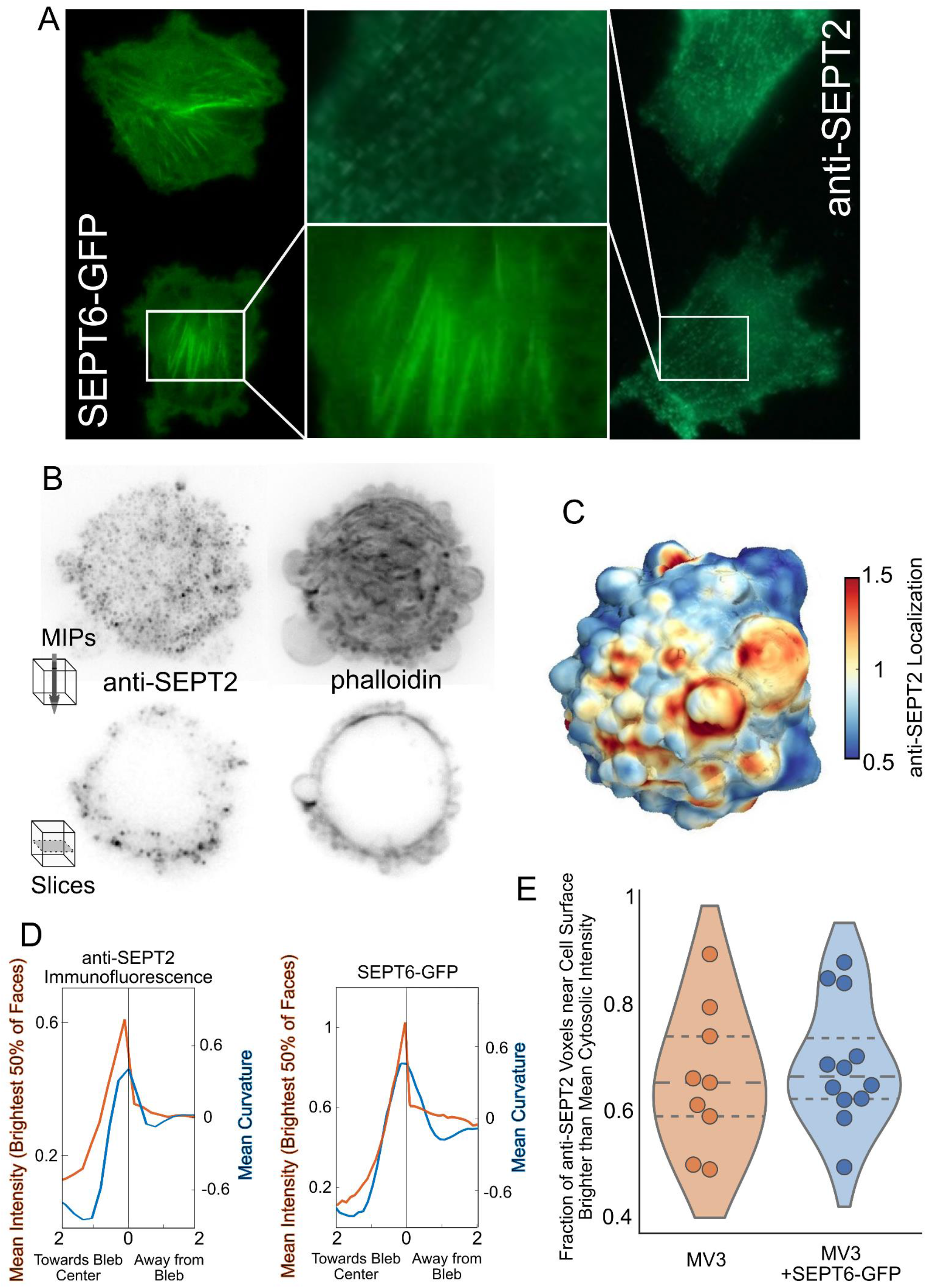
Endogenous septin localization via immunofluorescence. **(A)** MV3 cells adhered to fibronectin-coated glass slides showing either the murine SEPT6-GFP probe or anti-SEPT2 immunofluorescence localization. While the well-established septin localization to actin stress fibers (https://doi.org/10.1016/S1534-5807(02)00366-0) is apparent in both samples, the anti-SEPT2 signal is irregular, punctate, and possesses low signal-to-noise ratio (SNR) compared to SEPT6-GFP. **(B)** Anti-SEPT2 signal in a rounded blebbing MV3 cell embedded in soft bovine collagen, with maximum intensity projections (top) and single z-slices of 0.16 micron thickness (below). Phalloidin signal from the same cell shown on right to visualize the cell surface. Anti-SEPT2 signal possesses low SNR, as seen in adhered cells. Despite this, the signal is enriched at the cell surface in blebby regions of the cell just as seen in cells expressing the SEPT6-GFP probe (see **Fig S3**). **(C)** Cell surface distribution of anti-SEPT2 signal for the cell shown in **Fig S5B. (D)** Local anti-SEPT2 or SEPT6-GFP intensity and intracellular mean curvature as a function of distance from bleb edges. Mean intensity is comprised only of the brightest 50% of the cell surface to account for punctate and discontinuous immunofluorescent signal. **(E)** Fraction of cortical voxels (within 0.96 µm of surface) with anti-SEPT2 intensity higher than cytoplasmic mean intensity in MV3 cells with and without expression of SEPT6-GFP. Dashed lines separate quartiles and dots represent individual cells.

**Figure S7:**
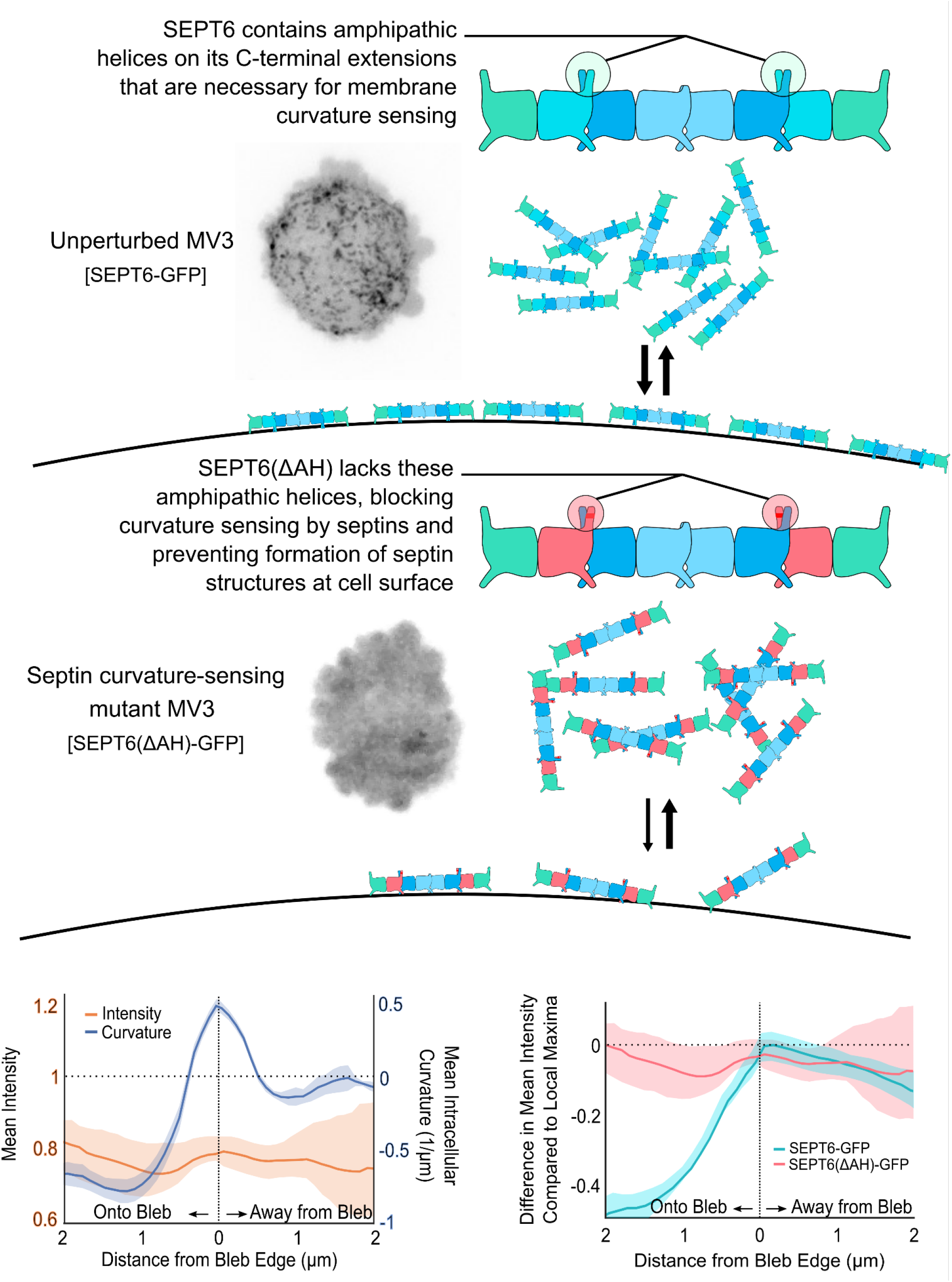
A mutation that disrupts curvature sensing blocks septin localization to the cell surface. Top, cartoon illustrating the mechanism underlying the altered function of the SEPT6(ΔAH)-GFP mutant. Bottom left, local SEPT6(ΔAH)-GFP intensity and intracellular mean curvature as a function of distance from bleb edges, calculated from 630 blebs across 5 MV3 cells. Error bands indicate 95% confidence intervals. Bottom right, difference in septin intensity compared to local maxima as a function of distance from bleb edge for both SEPT6-GFP (as seen in **Fig 2E**) and SEPT6(ΔAH)-GFP.

**Figure S8:**
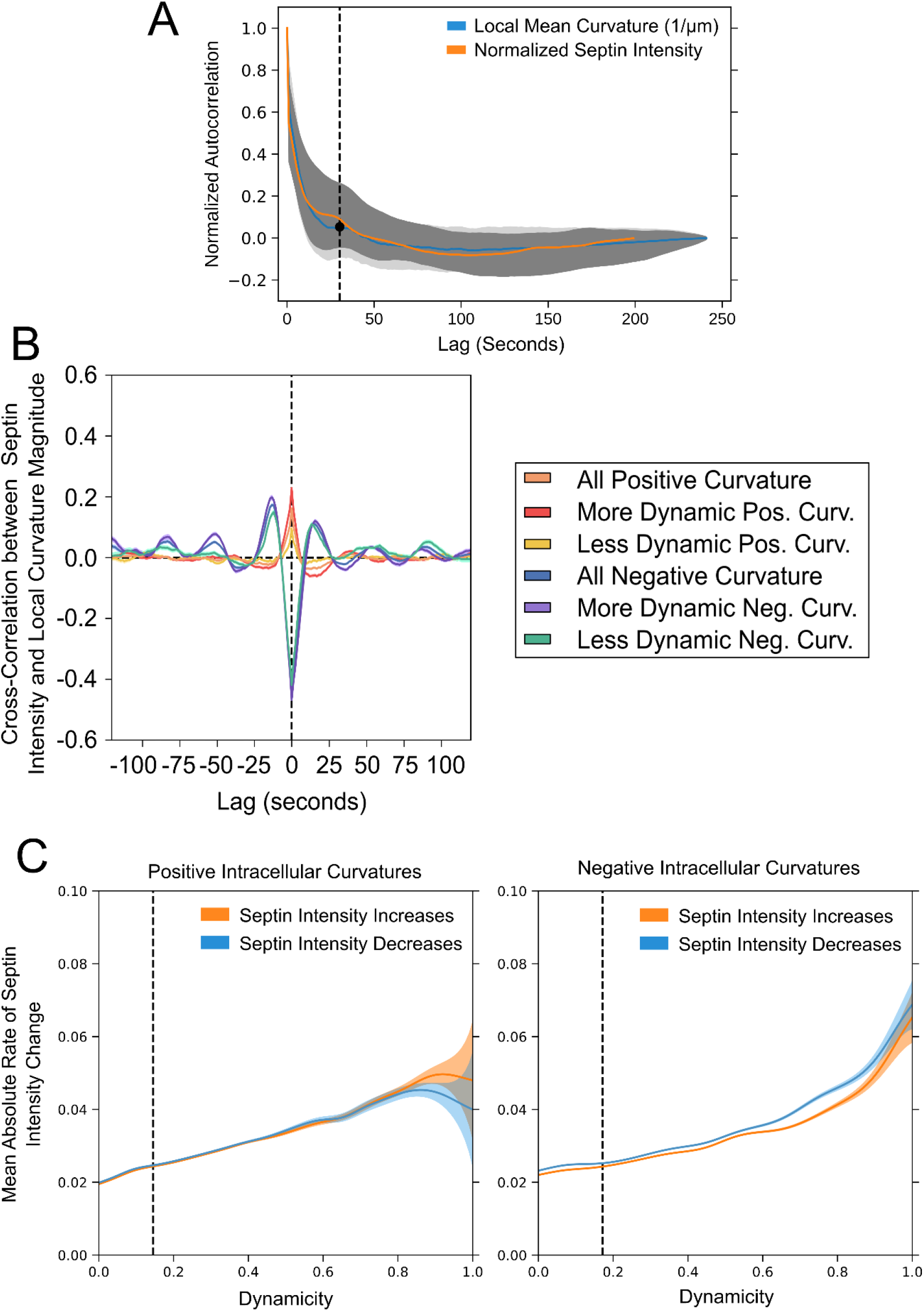
Analysis of high-speed timelapse SEPT6-GFP data in MV3. **(A)** Autocorrelation curves for the local mean curvature and SEPT6-GFP data presented in **Fig 2L**; the fluorescence signal was normalized to account for photobleaching as described in Methods... **(B)** Temporal cross-correlation functions as shown in **Fig 2L**, here with extended range of time lags. **(C)** Mean rates of absolute change in SEPT6-GFP intensity for positive and negative intracellular mean curvature contigs, expressed as a function of the dynamicity of the contigs they occur within. Dynamicity is defined as the total number of timepoints showing intensity increases and decreases that occur within a contig, divided by the total number of timepoints within that contig. Derived from the same dataset used in **Fig 2L**. Vertical dashed line represents the threshold for low and high dynamic groups as shown in **Figs 2L** and **S7B**.

**Figure S9:**
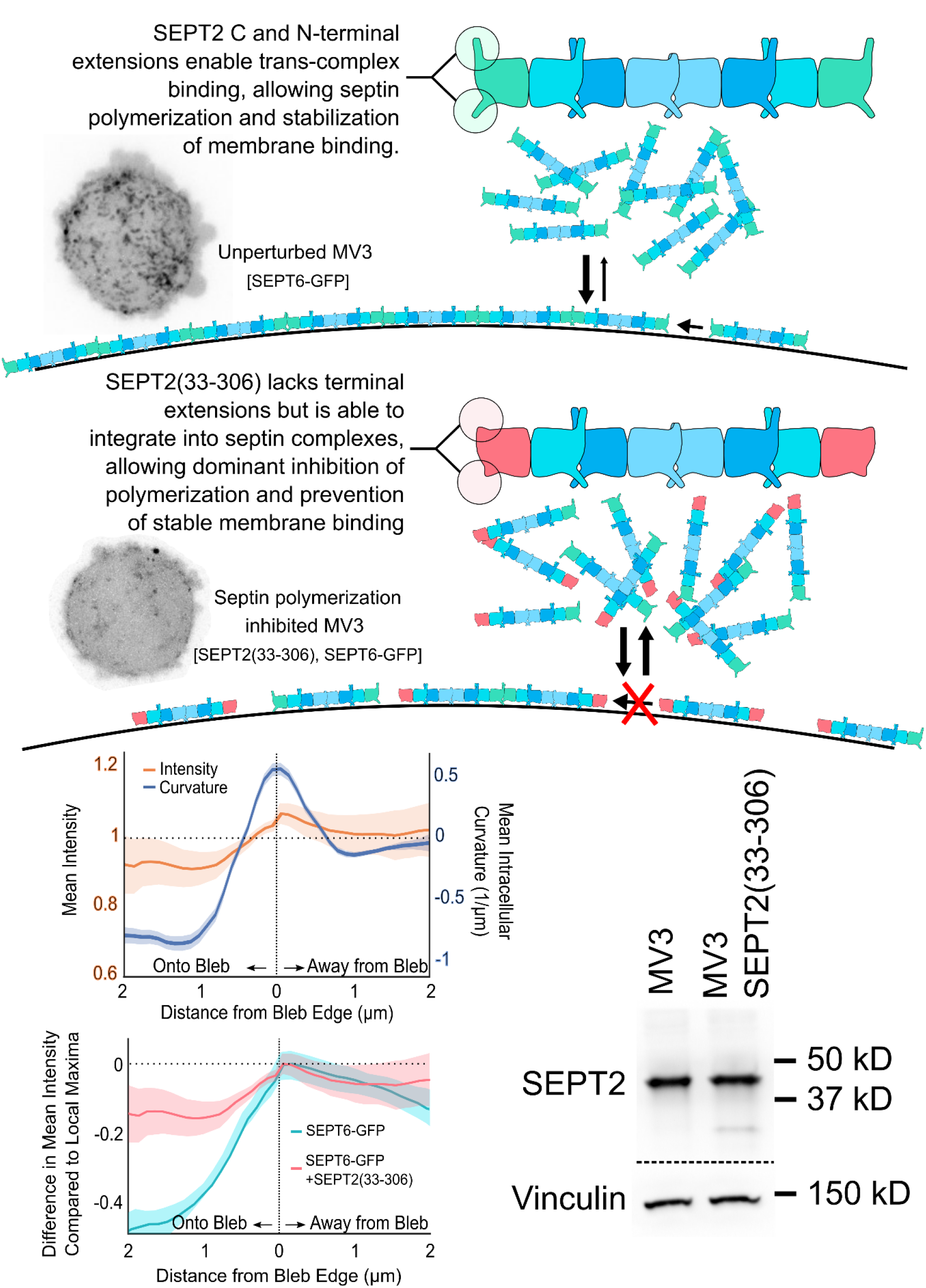
A mutation that disrupts inter-oligomeric polymerization blocks septin localization to the cell surface. Top, a cartoon illustrating the mechanism underlying the altered function of the SEPT2(33-306) mutant. Bottom left above, local SEPT6-GFP intensity and intracellular mean curvature as a function of distance from bleb edges in cells expressing SEPT2(33-306), calculated from 1293 blebs across 8 MV3 cells. Error bands indicate 95% confidence intervals. Bottom left below, difference in SEPT6-GFP intensity compared to local maxima as a function of distance from bleb edge for cells expressing SEPT2(33-306) and control cells (as seen in **Fig 2E**). Bottom right, Western blot of SEPT2 in MV3 cells with and without SEPT2(33-306) expression. Calculated mass of SEPT2 is 41.5kD and SEPT2(33-306) is 31.7 kD. Annotations show positions of the indicated ladder bands.

**Figure S10:**
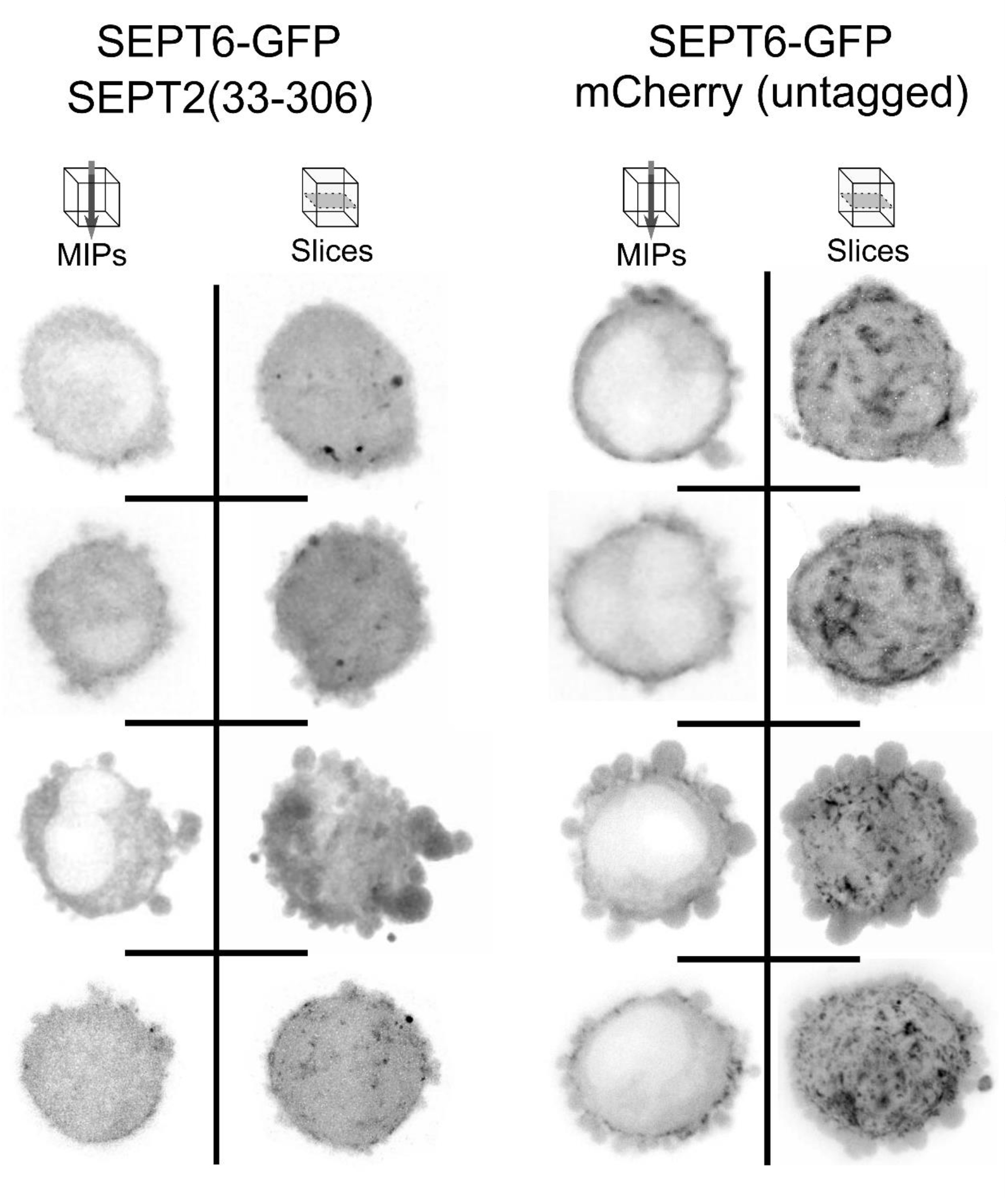
Effect of SEPT2(33-306) Expression on Septin Localization. Murine SEPT6-GFP probe localization in MV3 cells embedded in soft bovine collagen. Maximum intensity projections and single z-slices of 0.16 micron thickness shown for representative cells in two columns. (Left) Cells expressing the SEPT2(33-306) mutant. (Right) Cells expressing cytosolic mCherry (not shown) using the same pLVX-IRES-Hyg vector as the SEPT2(33-306) construct. Included as a negative control demonstrating that the septin mislocalization in SEPT2(33-306)-expressing cells is not due to effects arising from the protein expression construct.

**Figure S11:**
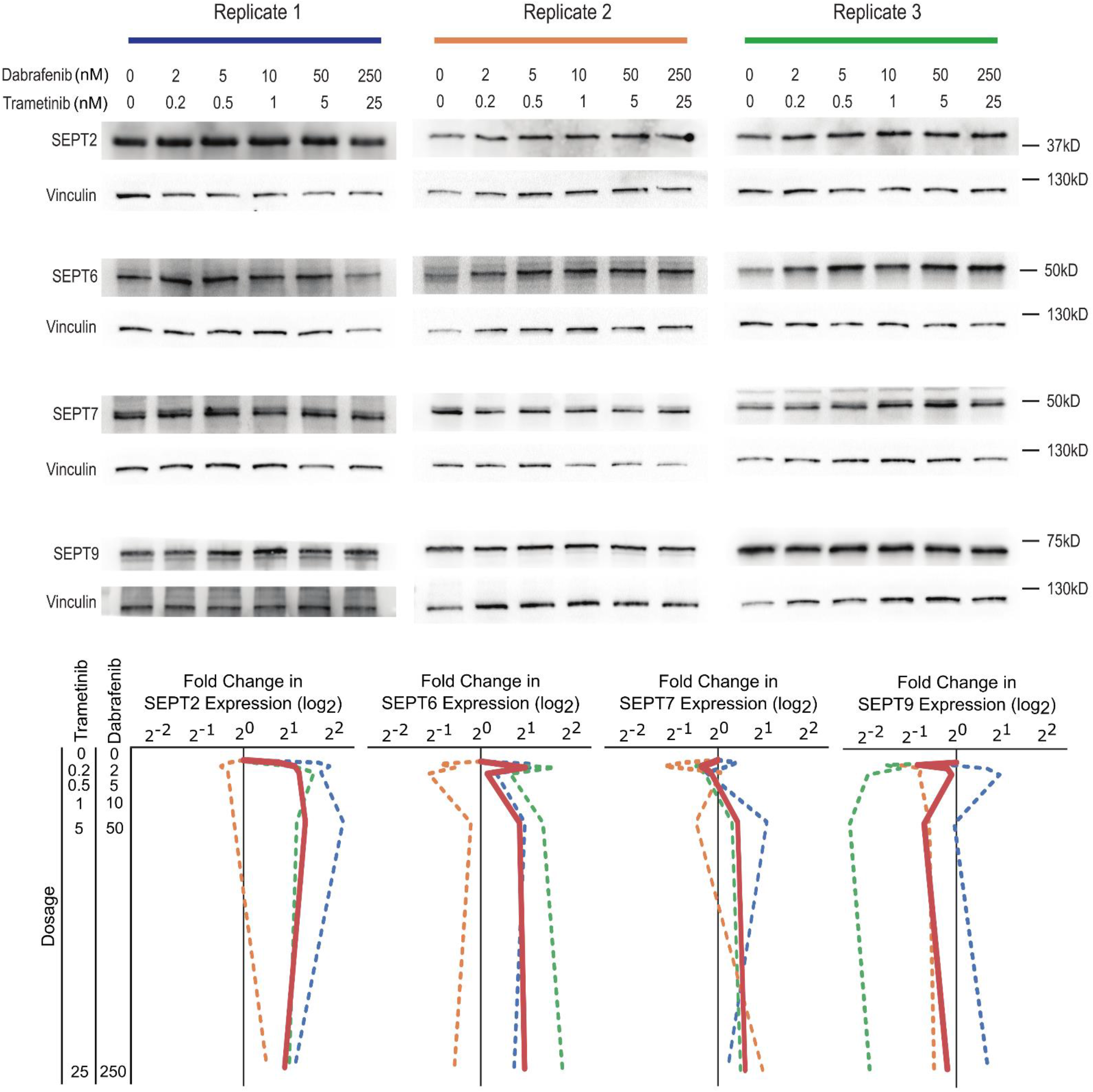
Changes in septin expression upon MAPKi treatment of A375 cells. Top, Western blots of SEPT2, SEPT6, SEPT7, and SEPT9 abundance (the 4 septins with highest expression in A375 cells, as measured by mass spectrometry, see **Table 2**) upon combined treatment with Dabrafenib and Trametinib at doses indicated. Three independent experiments were performed, with data from all replicates shown alongside vinculin loading controls. Bottom, densitometry analysis of Western blots showing fold change in septin expression as dose response curves. Dashed lines indicate individual experiments (color-coded according to the color bars indicating Western blot replicates); solid red lines indicate means of all replicates.

**Figure S12:**
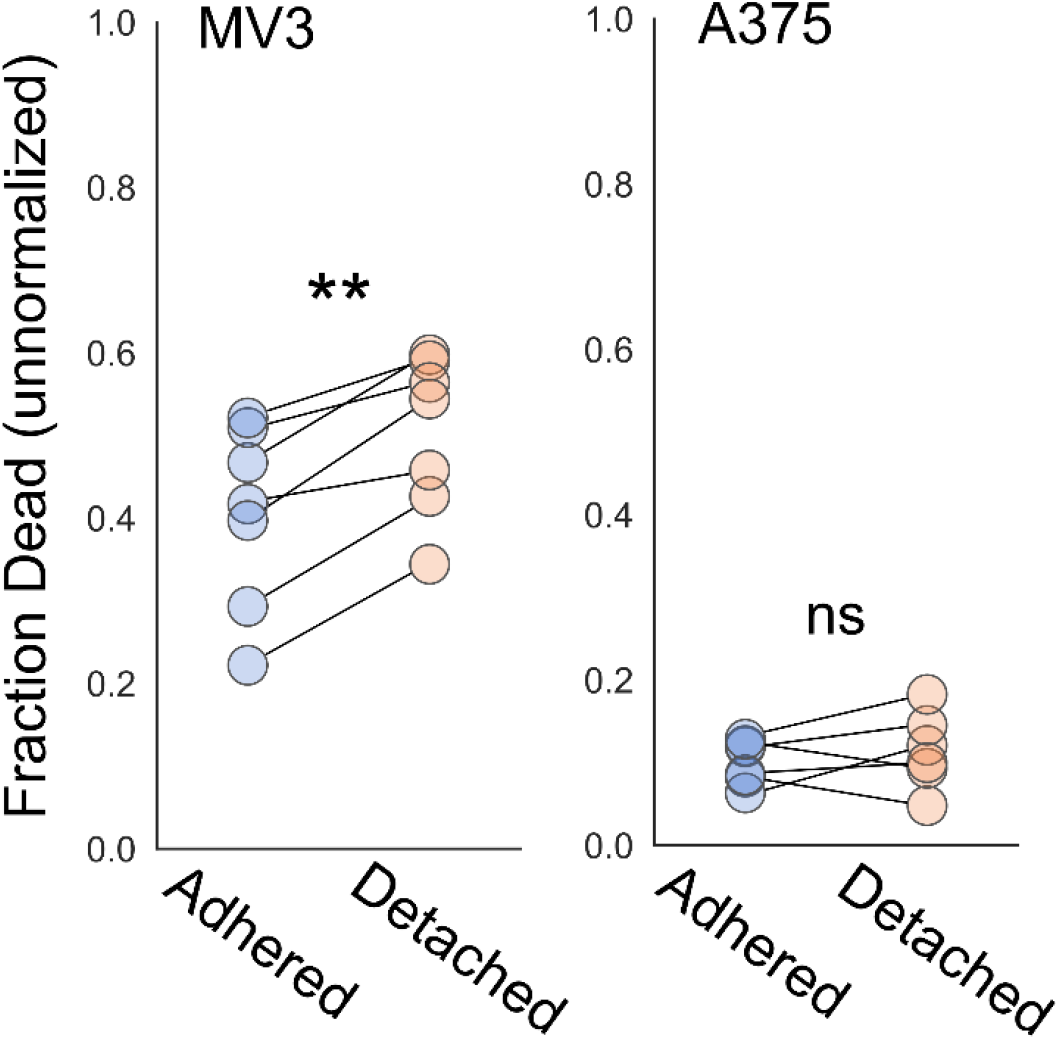
Viability effects of SEPT2(33-306) expression. Viability in adhered and detached melanoma cells expressing SEPT2(33-306). Cells grown for 24 hours and assayed for cell death using ethidium homodimer staining. All treatment groups grown and assayed in simultaneous paired experiments. Dots represent individual experiments. Cell counts for all replicates (see **Table 1** for individual counts): MV3 (951, 872), A375 (653, 1302). MV3 and A375 datasets were tested using Paired T Tests, p=0.001021 and p=0.3884

**Figure S13:**
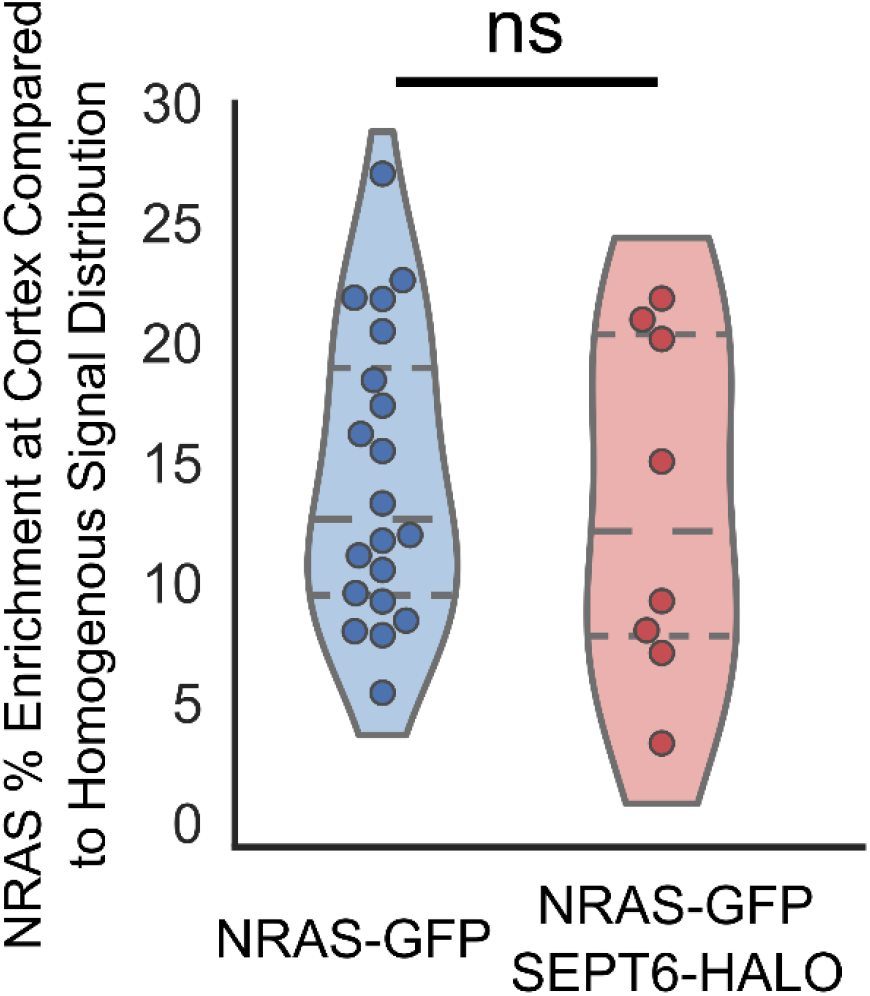
Effect of SEPT6 expression on NRAS Localization. Observed NRAS-GFP percent enrichment at the cortex (voxels within 0.96 µm of surface) of individual MV3 cells with and without expression of SEPT6-HALO using the same pLVX-IRES-Hyg vector as the SEPT2(33-306) construct. Included as a negative control demonstrating that the NRAS mislocalization in SEPT2(33-306)-expressing cells is not due to effects arising from the protein expression construct. Significance tested with two sample T-test with pooled variance (p<0.738), normality tested with Shapiro-Wilk (p=0.3152 & 0.4267), variance tested with two-tailed F test (p=0.181). Dashed lines separate quartiles.

**Figure S14:**
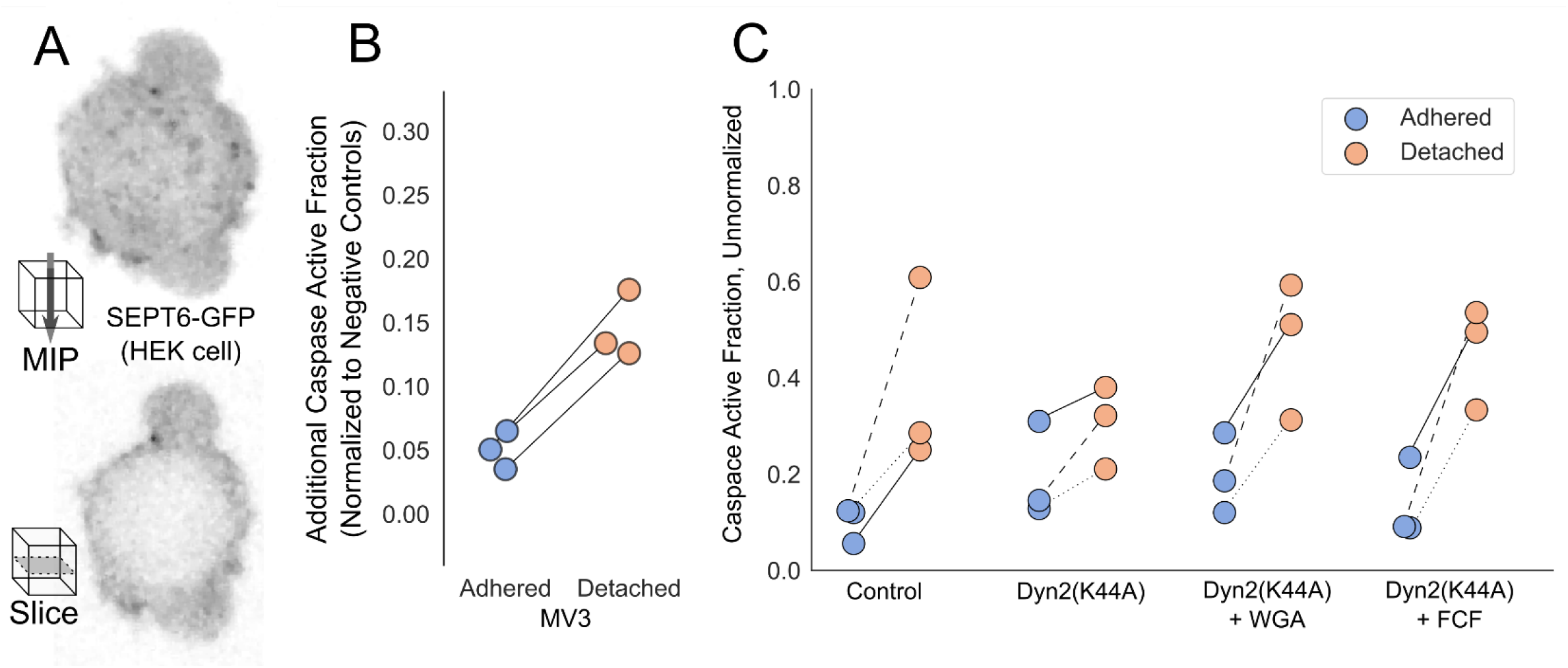
Anoikis resistance in non-malignant cells. **(A)** SEPT6-GFP localization in a recently detached HEK cell, embedded in soft bovine collagen. Top, maximum intensity projection (MIP); bottom, individual 0.16 µm z-slice from the same cell. **(B)** Additional fraction of MV3 cells showing caspase activation after 4 hours of treatment with 10 µg/mL WGA (compared to matched, same-day negative controls) for adhered and detached cells. Total cell counts with control in parenthesis (see **Table 1** for individual counts): adhered 459(529), detached 655(1026). **(C)** The same data shown in **Fig 5C**, without normalization. Solid, dashed, and dotted lines represent paired, same-day experiments. Total cell counts for each condition, with adhered group in parenthesis (see **Table 1** for individual counts): Control 164(168), DYN2(K44A) 187(178), DYN2(K44A)+WGA 160(183), DYN2(K44A)+FCF 196(137).

